# Macrophage extracellular traps promote maladaptive cardiac remodelling and heart failure via PAD4-dependent mechanisms

**DOI:** 10.64898/2026.03.15.711858

**Authors:** Shohei Ichimura, Tomofumi Misaka, Satoshi Okochi, Ryo Ogawara, Yu Sato, Shunsuke Miura, Tetsuro Yokokawa, Saori Miura, Koki Ueda, Masayoshi Oikawa, Akiomi Yoshihisa, Kazuhiko Ikeda, Takafumi Ishida, Yasuchika Takeishi

## Abstract

**Aims:** The activation of inflammatory cells, particularly macrophages, plays a pivotal role in the pathogenesis of cardiac remodelling and heart failure. Emerging evidence indicates that extracellular traps released from inflammatory immune cells contribute to the progression of various pathologies. However, the clinical relevance and mechanistic role of macrophage extracellular traps (METs) in heart failure remain to be elucidated.

**Methods and Results:** Endomyocardial biopsy specimens from 69 patients with heart failure were analysed by fluorescent immunostaining to identify and quantify METs. The numbers of METs per myocardial tissue area in patients with heart failure showed a negative correlation with left ventricular (LV) ejection fraction and a positive correlation with LV end-diastolic diameter. Patients with higher MET counts had significantly lower event-free survival from the composite cardiac events. In a murine model of pressure overload by transverse aortic constriction (TAC), METs were most abundantly observed at 3 days post-TAC and remained detectable throughout the 4-week observation period. In vitro, time-dependent MET formation was induced by an intrinsic trigger of mitochondrial DNA in bone marrow-derived macrophages from wild-type (WT) mice, but not in peptidyl arginine deiminase 4 (PAD4)-deficient macrophages, indicating that PAD4 activity is indispensable for MET formation. The recipient mice transplanted with bone marrow cells from PAD4 knockout mice showed more preserved cardiac function, reduced myocardial fibrosis, and improved survival in response to TAC, compared to those transplanted with WT mice. Ex vivo analyses demonstrated that conditioned medium containing METs from WT macrophages induced fibroblast-to-myofibroblast transition via Toll-like receptor 4 signalling.

**Conclusions:** PAD4-dependent MET formation from bone marrow-derived macrophages represents a novel driver of cardiac remodelling. Targeting MET formation may offer a potential therapeutic strategy for heart failure.

**Translational Perspective:** Macrophage extracellular traps (METs) are abundant in myocardial tissue from patients with heart failure with reduced ejection fraction and are associated with adverse left ventricular remodelling and worse clinical outcomes. These findings support myocardial MET burden as a potential tissue biomarker to improve risk stratification in heart failure patients. In mice, pressure overload induces MET formation, and hematopoietic PAD4 deficiency suppresses myocardial METs, attenuates fibrosis, preserves cardiac function, and improves survival. Mechanistically, mitochondrial DNA-enriched cardiomyocyte-derived exophers trigger PAD4-dependent METs, which activate cardiac fibroblasts through TLR4 signalling. Suppressing METs represents a potential therapeutic strategy to attenuate the progression of heart failure.

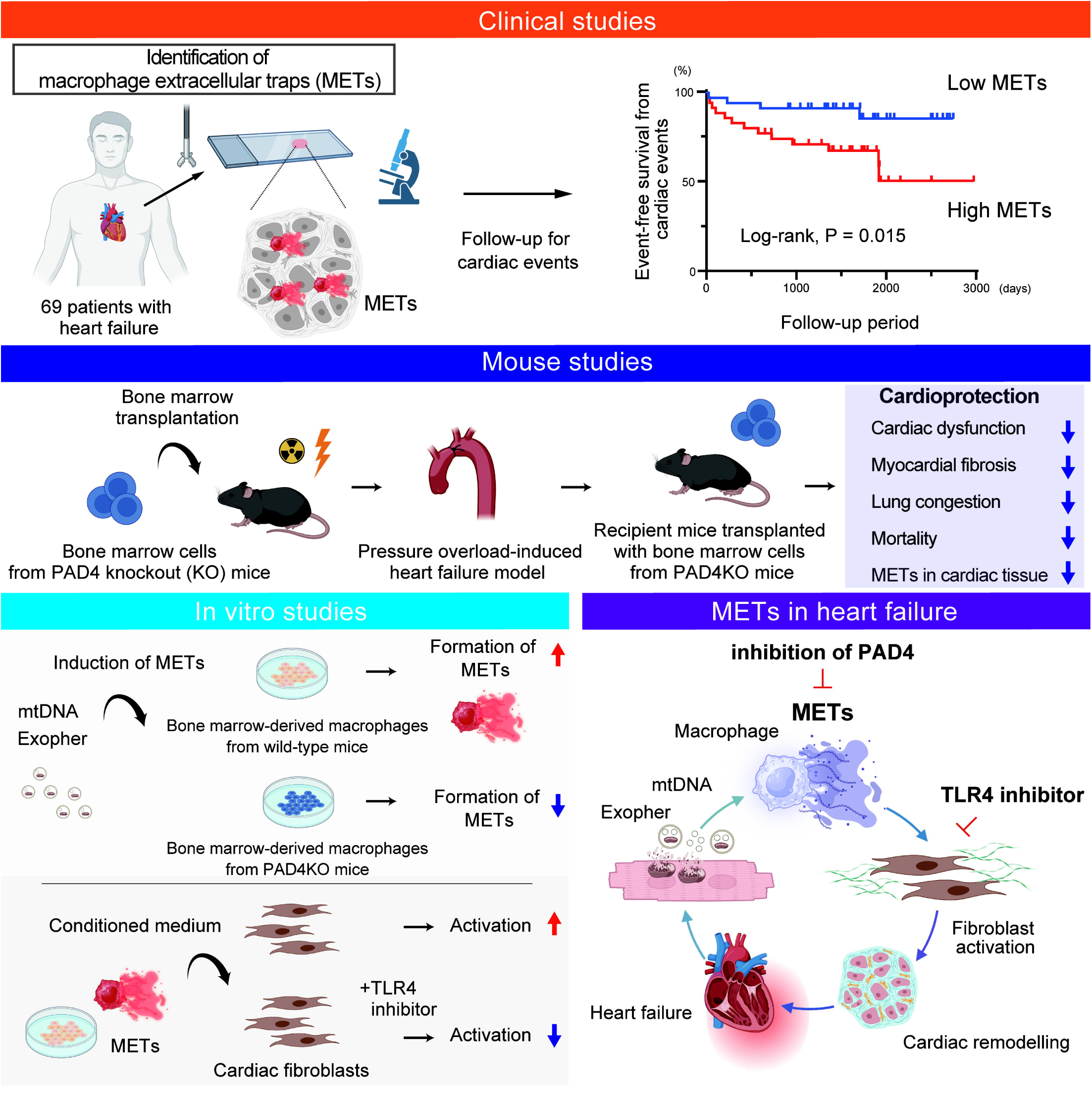

## **1.** Introduction

Heart failure develops through a sustained remodelling process in which the left ventricle undergoes hypertrophy, fibrotic expansion, and progressive dilatation.^1^ These structural alterations are closely linked to persistent inflammatory signalling within the myocardium, where recruited immune cells interact with resident cardiac cells and amplify tissue injury.^2^ Although inflammation has long been recognized as a hallmark of heart failure, immunosuppressive therapies have not translated into clear clinical benefit, underscoring the need to identify disease-promoting inflammatory mechanisms with greater precision.^3^

Among infiltrating myeloid populations, bone marrow-derived macrophages (BMDMs) are regarded as central drivers of myocardial inflammation and adverse remodelling.^4,5^ Yet, the precise mechanisms by which macrophages drive maladaptive remodelling in the failing heart remain incompletely understood.

It is increasingly recognized that inflammatory leukocytes undergo ETosis, releasing extracellular traps composed of DNA, histones, and granule-derived proteins.^6,7^ In particular, neutrophil extracellular traps have been implicated in sterile inflammation and cardiovascular diseases.^8,9^ More recently, macrophages have also been shown to generate extracellular traps, termed as macrophage extracellular traps (METs).^10^ However, whether METs are present in failing cardiac tissue and whether METs contribute to the pathogenesis of heart failure remain unclear.

Here, the present study investigated the clinical and pathological significance of METs in heart failure and tested the hypothesis that inhibition of MET formation may be a potential approach to prevent the development of heart failure.

## 2. Methods

Detailed methods are available in Supplementary data online. The data that support the findings of this study are available from the corresponding author upon reasonable request.

### 2.1 Patient cohort

Sixty-nine patients with symptomatic stage C or D heart failure with reduced ejection fraction (HFrEF) due to idiopathic dilated cardiomyopathy, who were admitted to Fukushima Medical University Hospital and underwent endomyocardial biopsy between April 2016 and March 2021, were enrolled in this study.^11^ Biopsy samples were collected from the interventricular septum using a right ventricular approach.^12^ The control group included age- and sex-matched subjects without a history of heart failure. Although they were evaluated for suspected cardiac disease, no specific diagnosis was identified, including on endomyocardial biopsy. The study protocol was approved by the Ethics Committee of Fukushima Medical University (approval numbers, 823, 2020-309, and REC2024-161). The investigation conformed to the principles outlined in the Declaration of Helsinki, and all patients provided written informed consent prior to study enrollment.

### 2.2 Fluorescent immunohistochemistry on the myocardial biopsy specimen

Fluorescent immunohistochemistry was carried out on paraffin-embedded myocardial biopsy sections to identify METs. After deparaffinization and rehydration, the 2-μm-thick sections were stained with the following primary antibodies: citrullinated histone H3 (1:100, ab5103, Abcam), CD68 (1:100, ab955, Abcam), troponin I (1:100, ab188877, Abcam) followed by the appropriate secondary antibodies, including Donkey anti-Rabbit IgG (H+L) Highly Cross-Adsorbed Secondary Antibody, Alexa Fluor 647 (A31573, Thermo Fisher Scientific), Donkey anti-Mouse IgG (H+L) Highly Cross-Adsorbed Secondary Antibody, Alexa Fluor 594 (A21203, Thermo Fisher Scientific) or Donkey Anti-Goat IgG H&L, Alexa Fluor 488 (ab150129, Abcam), and then mounted with DAPI-containing mounting media (Fluoro-Gel II with DAPI, Electron Microscopy Sciences). All microscopic images were obtained with a Keyence BZ-X700 microscope and processed with the BZ II Viewer software (Keyence). For quantitative analysis, imaging was performed systematically across the entire field of each human biopsy sample, independently by investigators blinded to the patients’ clinical characteristics and outcomes.

### 2.3 Clinical data and patient outcomes

Clinical characteristics were documented in accordance with our standard clinical practice, including demographic information, laboratory findings, echocardiographic parameters, right heart catheterization–derived hemodynamics, cardiac magnetic resonance imaging, and cardiopulmonary exercise testing.^8^ After hospital discharge, patients were followed through review of medical records from our institution and affiliated referral centers to document cardiac events. The primary outcome was a composite of major cardiac events, comprising cardiac death, worsening heart failure, and implantation of a left ventricular assist device (LVAD) as a bridge to transplantation. Cardiac death was defined as death resulting from progressive heart failure, ventricular arrhythmias such as ventricular tachycardia or ventricular fibrillation, or sudden cardiac death. Worsening heart failure was defined as the development of clinical signs and symptoms requiring urgent intervention and subsequent hospitalization. All clinical endpoints were adjudicated by at least two independent cardiologists.

### 2.4 Animals

B6.Cg-Padi4tm1.1Kmow/J (PAD4 KO, stock number #030315) and B6.Cg-Gt(ROSA)26Sortm14(CAG-tdTomato)Hze/J (tdTomato, stock number #007914) were obtained from Jackson Laboratory. Transgenic mice expressing α-myosin heavy chain promoter-driven Cre recombinase were described,^13^ and crossed with tdTomato mice to generate cardiomyocyte-specific tdTomato-expressing mice. CAG-EGFP reporter mice were purchased from Japan SLC. C57BL/6 wild-type (WT) male mice were obtained from Japan SLC or Jackson Laboratory Japan. Mice were housed with food and water ad libitum during 12-h light/12-h dark cycles (light, 7:00–19:00; dark, 19:00–7:00), and ambient temperature (21.5 °C) and humidity (55 ± 10%) were monitored. All experiments were performed in 8- to 14-week-old male mice in a C57BL/6 background. All animal studies were approved by Fukushima Medical University Animal Research Committee (approval numbers 2021016 and 2023083) and were performed in accordance with the Guide for the Care and Use of Laboratory Animals and institutional guidelines for animal care.

### 2.5 Bone marrow transplantation (BMT)

Recipient male C57BL/6J mice aged 8–10 weeks were lethally irradiated with a total dose of 9.0 Gy using an experimental X-irradiation device (MBR-1605R, Hitachi Power Solutions Co., Ltd.) 24 h prior to BMT.^14^ Whole bone marrow cells were collected from donor femurs and tibiae, washed with PBS, and 5.0 × 10^6^ cells were injected into recipient mice via the tail vein.

### 2.6 Animal model of pressure overload by transverse aortic constriction (TAC)

Mice were subjected to TAC or sham surgeries as previously reported.^15,16^ Mice were anesthetized with a single intraperitoneal administration of medetomidine (0.3 mg/kg), midazolam (4.0 mg/kg), and butorphanol (5.0 mg/kg). Adequacy of anaesthesia was confirmed by the absence of pedal withdrawal and corneal reflexes, together with a stable respiratory pattern. Once a surgical depth of anaesthesia was achieved, a trans-sternal thoracotomy was performed. TAC procedure was conducted by placing a 7-0 suture around the aortic arch between the brachiocephalic artery and the left common carotid artery, using a 27-gauge blunt needle to standardize the degree of narrowing. After removal of the needle, a calibrated stenosis remained. Sham-operated mice underwent the same procedure without ligation of the aorta. Following chest closure, the animals were monitored until full recovery. At the experimental endpoint, mice were euthanized under deep anaesthesia with isoflurane followed by cervical dislocation.

### 2.7 Echocardiographic assessment in mice

Transthoracic echocardiography was performed with a Vevo 2100 high-resolution imaging system (VisualSonics Inc.) using either a 20- or 40-MHz transducer.^15^ Parasternal short-axis M-mode recordings at the level of the papillary muscles were acquired in conscious mice. LV fractional shortening was calculated as 100 × (LVIDd − LVIDs) / LVIDd, where LVIDd and LVIDs denote end-diastolic and end-systolic LV internal dimensions, respectively. The values were calculated as the mean of at least four consecutive cardiac cycles. The peak pressure gradient across the aortic arch was assessed by pulsed-wave Doppler in mice lightly anesthetized with isoflurane (0.5–1.5%), titrated to maintain a heart rate of approximately 400 beats per minute.

### 2.8 Histological analysis and fluorescent immunohistochemistry of murine hearts

Murine LV samples were fixed in 4% paraformaldehyde solution for paraffin embedding or embedded in the O.C.T. compound (Tissue-Tek). Paraffin-embedded sections (2 um thickness) were stained with hematoxylin-eosin or Elastica-Masson for assessment of myocardial fibrosis.^15^ Myocardial fibrosis fractions were measured using ImageJ software (National Institutes of Health). For the assessment of cardiomyocyte cross-sectional area, paraffin-embedded tissue sections were incubated with wheat germ agglutinin conjugated to Alexa Fluor 488 (Thermo Fisher Scientific) and mounted with DAPI-containing medium.^15^ Cardiomyocyte cross-sectional area was quantified from more than 100 cells using ImageJ software. For immunohistochemical staining, the paraffin-embedded tissue sections were incubated with the primary antibody of alpha-smooth muscle actin (19245, Cell Signaling Technology) followed by anti-rabbit IgG antibody labeled with peroxidase (414341, Nichirei Bioscience) with DAB peroxidase substrate system (347-00904, Dojin Co., Ltd.) and counterstained with hematoxylin. Fluorescent immunohistochemistry was carried out on O.C.T. compound-embedded LV sections (6 um thickness). The sections were incubated with primary antibodies against citrullinated histone H3 (ab5103, Abcam), troponin I (ab188877, Abcam), CD68 (MCA1957, Bio-Rad) or neutrophil elastase (MAB4517, R&D Systems), followed by appropriate secondary antibodies, including Donkey Anti-Rat IgG H&L, Alexa Fluor 594 (ab150156, Abcam), Donkey anti-Goat IgG H&L, Alexa Fluor 488 (ab150129, Abcam), Donkey anti-Rabbit IgG (H+L) Highly Cross-Adsorbed Secondary Antibody, Alexa Fluor 647 (A31573, Thermo Fisher Scientific) or Donkey Anti-Rabbit IgG H&L Alexa Fluor 750 (ab175731, Abcam), and then mounted with a DAPI-containing mounting medium. All microscopic images were obtained with a Keyence BZ-X700 microscope and processed with the BZ II Viewer software.

### 2.9 Western blot analysis

Snap-frozen mouse heart samples or cultured cells were initially homogenized in lysis buffer (Cell Lysis Buffer, Cell Signaling Technology) containing protease inhibitor cocktail (BD Biosciences). Protein concentrations were quantified using the BCA Protein Assay Kit (Pierce, Thermo Fisher Scientific). Equal amounts of protein were resolved by SDS-polyacrylamide gel electrophoresis and transferred to nitrocellulose membranes (Amersham Protran Premium 0.2 NC, GE Healthcare). After the membranes were blocked using 5% bovine serum albumin or 5% skim milk for 1 h, membranes were incubated overnight at 4°C with the following specific antibodies: BNP (ab239510, Abcam), Collagen III (GTX111643, GeneTex), GAPDH (60004-1-Ig, Proteintech), PAD4 (ab21480, Abcam), and Histone H3 (9715, Cell Signaling Technology). After incubation with the appropriate IRDye 680 or IRDye 800 secondary antibodies (926-68070, 926-68071, 926-32210; LI-COR Biosciences), fluorescent signals were visualized using an Odyssey CLX imaging system (LI-COR Biosciences) and band intensities were quantified using ImageJ software (NIH) or Image Studio software (LI-COR Biosciences).

### 2.10 Cell culture, preparation for BMDMs, and magnetic-activated cell sorting

RAW264.7 cells, a murine macrophage cell line, were purchased from KAC Co., Ltd. and cultured in DMEM (Fujifilm Wako) supplemented with 10% FBS (Gibco), 100 U/mL penicillin and 100 μg/mL streptomycin (Fujifilm Wako) at 37 °C in a humidified atmosphere containing 5% CO_2_. GSK484, PAD4 inhibitor was purchased from Abcam (ab223598). BMDMs were generated by culturing mouse bone marrow cells from WT mice or PAD4KO mice in DMEM supplemented with 10% FBS, 100 U/mL penicillin, 100 μg/mL streptomycin and 10 ng/mL recombinant M-CSF (R&D Systems).^17^ The culture medium was replaced every two days for a total of 7 days. Magnetic-activated cell sorting was performed on bone marrow cells using MACS MS columns (Miltenyi Biotec GmbH) with CD11b MicroBeads (130-097-142, Miltenyi Biotec GmbH) according to the manufacturer’s instructions.

### 2.11 Immunofluorescence and time-lapse live-cell imaging

Cells were fixed with 4% paraformaldehyde, permeabilized with either 0.05% or 0.2% Triton X-100 for 3 or 10 min, and subsequently blocked with 2% bovine serum albumin for 1 h at room temperature and incubated with specific primary antibodies: anti-citrullinated histone H3 (ab5103, Abcam), CD68 (MCA1957, Bio-Rad), followed by appropriate secondary antibodies such as Goat anti-Rabbit IgG (H+L) Highly Cross-Adsorbed Secondary Antibody, Alexa Fluor 647 (A21245, Thermo Fisher Scientific), Anti-rat IgG (H+L), Alexa Fluor 488 (4416, Cell Signaling Technology) and then mounted with DAPI. For time-lapse live-cell imaging, BMDMs were seeded on 35-mm μ-ibiTreat high imaging dishes (Ibidi) 24 h before imaging. Cells were incubated with SiR-DNA (30 min at 1 µM, Cytoskeleton, CY-SC007) and SYTOX Green (Thermo Fisher Scientific) according to the manufacturers’ protocols.

### 2.12 Administration of anti-Ly6G antibody in mice

Neutrophil depletion was induced by intraperitoneal injection of 100 µg anti-Ly6G monoclonal antibody (InVivoPlus, clone 1A8, BioXcell) administered twice weekly in mice. Control mice received an equivalent dose of isotype control antibody (rat IgG2a, clone 2A3, BioXcell).^18,19^

### 2.13 Flow cytometry of peripheral blood

Peripheral blood was collected from the tail vein into EDTA-2Na-containing tubes. Following red blood cell lysis with an ammonium chloride-based lysis buffer, leukocytes were stained with CD45.2-APC (109814, BioLegend), Gr-1-PE (108407, BioLegend), and CD11b-FITC (11-0112-82, Invitrogen). Flow cytometric analysis was performed using a Bigfoot Spectral Cell Sorter (Invitrogen, Thermo Fisher Scientific). The gating strategies are provided in Supplementary data online.

### 2.14 Sorting exophers from cardiac tissue

Freshly excised mouse hearts were digested with 1Lmg/mL collagenase type II (Worthington Biochemical) and 1 U/ml DNase I (Sigma-Aldrich) for 45 min at 37°C.^20^ The resulting single-cell suspension was sequentially centrifuged at 100 g, 300 g, and 1000 g. The final pellet, enriched in cardiac exophers, was resuspended and subsequently analysed by flow cytometry (BD FACSMelody Cell Sorter, BD Biosciences, or Invitrogen Bigfoot Spectral Cell Sorter, Thermo Fisher Scientific), and data were analysed with FlowJo (version 10.10, Tree Star Inc.)

### 2.15 Transmission electron microscopy

The morphology of isolated exophers was examined by transmission electron microscopy using a negative staining method with 2% phosphotungstic acid (pH 7.0). Imaging was performed following standard protocols at Tokai Electron Microscopy, Inc. using a JEM-1400Plus (JEOL Ltd.) operated at 100 kV.

### 2.16 Mitochondrial DNA quantification

DNA was extracted and purified from mouse heart tissue using the QIAamp DNA Mini Kit (Qiagen) in accordance with the manufacturer’s instructions, including RNase A treatment. Quantitative PCR was performed to quantify mitochondrial DNA using THUNDERBIRD SYBR qPCR Mix (Toyobo Co., Ltd.) in a CFX Connect real-time PCR System (Bio-Rad) with Bio-Rad CFX Manager 3.1 software (Bio-Rad). Primers targeting the Co1 gene were used, while nuclear DNA was amplified with primers for the Ndufv1 gene. The primer sequences were as follows: Co1 forward 5′-TGCTAGCCGCAGGCATTAC-3′ and reverse 5′-GGGTGCCCAAAGAATCAGAAC-3′; Ndufv1 forward 5′-CTTCCCCACTGGCCTCAAG-3′ and reverse 5′-CCAAAACCCAGTGATCCAGC-3′. All reactions were performed in duplicate. Relative mitochondrial DNA abundance was determined by the ΔCT method, and values were normalized to the control group and expressed as fold changes.^16^

### 2.17 Liquid chromatography-tandem mass spectrometry (LC-MS/MS) and proteomics analysis

The LC-MS/MS was performed using an Easy-nLC1000 system (Thermo Fisher Scientific) coupled to an Orbitrap Elitemass spectrometer (Thermo Fisher Scientific) as previously described.^8^ Experimental triplicate runs were performed. The data from our prior study were used as reference data for comparison.^8^ Data were exported from Scaffold 5 (Proteome Software, Inc).

### 2.18 Isolation and culture of primary cardiac fibroblasts

Primary cardiac fibroblasts were isolated from mouse LVs. The LVs were minced into small pieces and digested using collagenase type 2 (1 mg/mL), and the collected cells were cultured in DMEM with 20% FBS. Prior to experiments, the cells were serum-starved for 24 h in DMEM containing 1% FBS. TLR4 inhibitor (TLR4-IN-C34, Selleck) was administered 1h prior to conditioned medium from BMDMs. Cells were fixed and immunofluorescent analysis was conducted with anti-alpha-smooth muscle actin antibody (19245, Cell Signaling Technology) and wheat germ agglutinin (Alexa Fluor 488 conjugated, Thermo Fisher Scientific).

### 2.19 Statistical analysis

Comparisons between two groups were conducted using either the unpaired Student’s t-test, or the Mann–Whitney U-test when appropriate. For analyses involving more than two groups, one-way ANOVA followed by Tukey’s post hoc test for multiple comparisons, or two-way repeated-measures analysis of variance followed by Sidak’s multiple comparisons test was applied. Categorical variables were assessed using Fisher’s exact test or the Chi-square test. Correlations were assessed using the Spearman correlation analysis. Receiver operating characteristic (ROC) curves were used to determine the optimal cutoff points for the primary composite cardiac outcomes and the area under the curve was calculated. Kaplan–Meier method was used for survival analysis. Univariate and multivariate Cox regression were used to analyze the independent predictors of prognosis. Statistical analyses were carried out using SPSS software version 26 (IBM Corp.) or GraphPad Prism version 8.1.2 (GraphPad Software). A P value < 0.05 was considered statistically significant.

## 3. Results

### 3.1 METs identified in the myocardial biopsy specimens from patients with heart failure and their association with adverse clinical outcomes

We first sought to identify the presence of METs and their clinical significance in patients with HFrEF. We employed histological approaches using myocardial biopsy specimens from HFrEF patients diagnosed as idiopathic dilated cardiomyopathy. METs were determined by identifying the cell structure of citrullinated histone H3, an essential and specific marker for the formation of ETosis^10^ and CD68, a marker for macrophages, along with DAPI-stained nuclei but negative for troponin I (Figure 1A). Clinical characteristics of patients (n=69) are summarized in Supplementary Table S1. The mean age was 55.5 ± 13.0 years with 84.1% being male and left ventricular ejection fraction (LVEF) of 26.9 ± 8.3%, and B-type natriuretic peptide (BNP) of 178.8 (89.2-379.1) pg/mL. Histogram distribution of MET counts per myocardial tissue area and the MET-to-macrophage ratio is presented in Figure 1B. The numbers of METs were positively correlated with those of CD68 macrophages as well as with the MET-to-macrophage ratio (Figure 1C).

**Figure 1.**
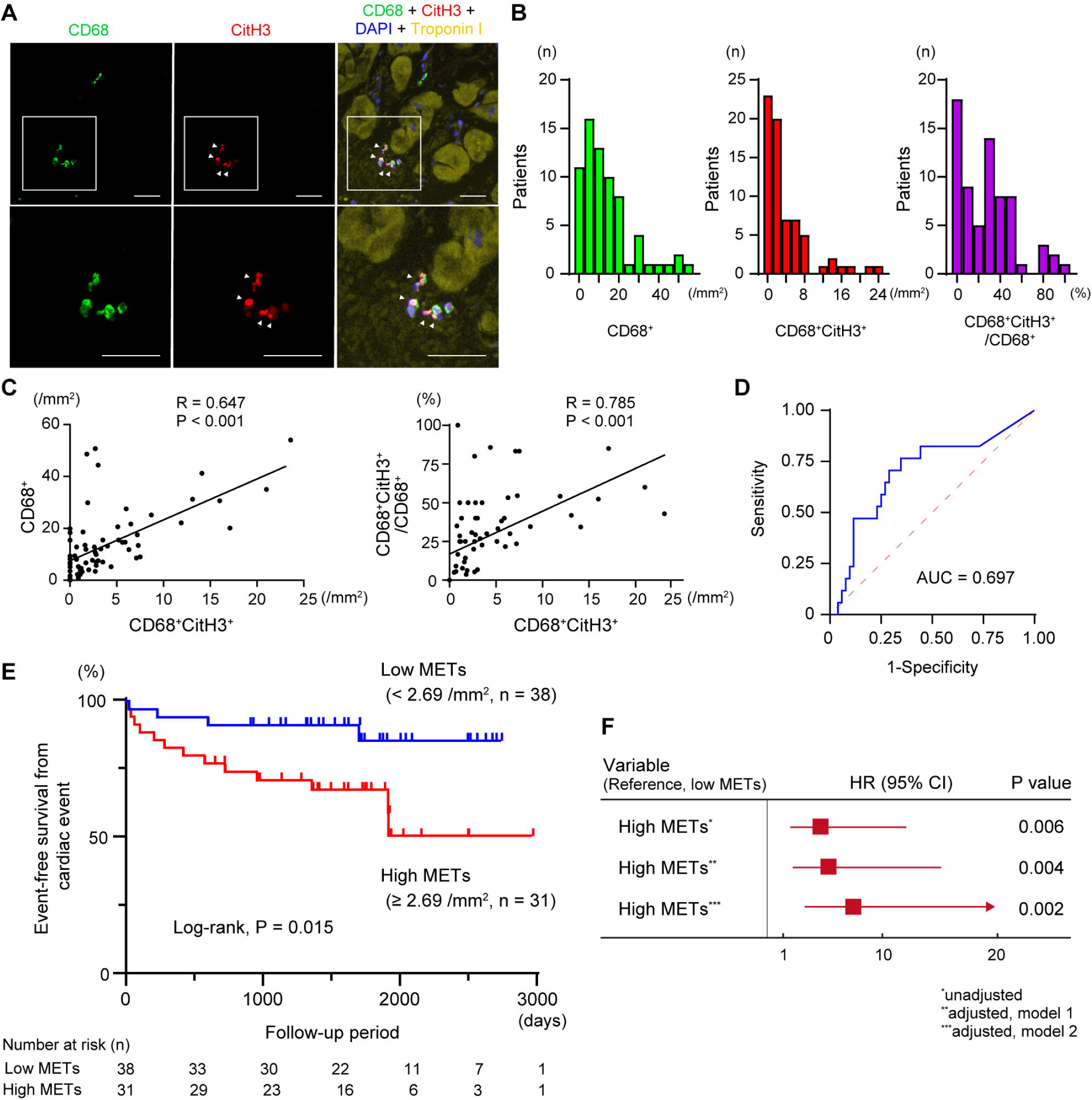
Identification of macrophage extracellular traps (METs) in myocardial tissue in patients with heart failure and their associations with clinical outcomes. **A**, Representative fluorescent immunohistochemistry images of myocardial biopsy specimens showing the identification of METs using antibodies against CD68 (green), citrullinated histone H3 (CitH3, red), and troponin I (yellow), along with nuclear staining using DAPI (blue). METs are defined as structures positive for CD68, CitH3, and DAPI, but negative for troponin I. Images at higher magnification are displayed in the lower panels from the boxed area in the upper panels. White arrowheads indicate MET-positive cells (CD68^+^CitH3^+^). Scale bars, 50 μm. **B**, Quantitative analyses illustrating the histogram distribution of the numbers of CD68^+^ macrophages and METs (CD68^+^CitH3^+^) per myocardial tissue area, and MET-to-macrophage ratio (CD68^+^CitH3^+^/CD68^+^) in patients with heart failure (n=69). **C**, Correlation analysis between MET count and CD68^+^ macrophage number or the MET-to-macrophage ratio (CD68^+^CitH3^+^/CD68^+^). **D**, Receiver operating characteristic curve analysis of myocardial MET counts (CD68^+^CitH3^+^) for predicting cardiac events including cardiac death, re-hospitalizations due to decompensated heart failure, and left ventricular assist device implantation. **E**, Kaplan-Meier analysis for the composite endpoint of cardiac events. Patients were stratified into low- and high-MET groups based on an optimal threshold of MET count of 2.69 per mm^2^ in myocardial tissue. The log-rank test was performed for the statistical comparison. **F**, Forest plot showing hazard ratios from univariate and multivariate Cox proportional hazards models for the composite cardiac outcome. Multivariate model 1 was adjusted for age and sex, and model 2 was adjusted for age, sex, body mass index, and left ventricular ejection fraction.

Compared to patients with non-heart failure controls (n=10), patients with heart failure showed significantly higher myocardial MET counts (Supplementary Figure S1). To assess the clinical relevance of MET burden, myocardial MET counts were positively associated with LV diameter or volume at both end-diastole and end-systole, and inversely correlated with LVEF, suggesting a link between METs and adverse LV remodelling in patients with heart failure (Supplementary Table S2).

Next, to evaluate the prognostic significance of myocardial METs, patients were followed for the primary composite outcome of cardiac death, worsening heart failure, or LV assist device implantation over a median of 1594 days. ROC curve analysis identified an optimal threshold of 2.69 METs/mm² for predicting cardiac events, with an area under the curve of 0.696 (95% confidence interval, 0.544–0.849; P=0.015, Figure 1D). When patients were stratified into two groups based on the threshold myocardial MET count, those with high MET counts showed significantly greater LV end-diastolic and end-systolic volumes (Supplementary Table S3). Kaplan–Meier analysis revealed that patients with high METs had lower event-free survival compared with those with low METs (Figure 1E). The univariable and multivariable Cox proportional hazards analyses further demonstrated that higher MET counts were associated with an increased risk of cardiac events (Figure 1F and Supplementary Table S4). In addition, myocardial MET counts analysed as a continuous variable were also associated with a higher risk of cardiac events (Supplementary Table S4). Taken together, these findings suggest that higher myocardial MET burden is associated with adverse outcomes in patients with HFrEF.

### 3.2 METs are induced by pressure overload in a murine model of HFrEF

We subsequently determined the dynamic changes in myocardial METs under stress-induced pathological conditions using a murine model of pressure overload by means of TAC.^16^ In this HFrEF model, TAC induced MET formation in LV tissue in WT mice at 1 day, 3 days, and 4 weeks after TAC (Figure 2A and 2B). Notably, METs were most abundantly observed at 3 days post-TAC and remained detectable throughout the 4-week observation period. These findings suggest that pathological stress caused by pressure overload induces sustained MET formation in the left ventricle.

**Figure 2.**
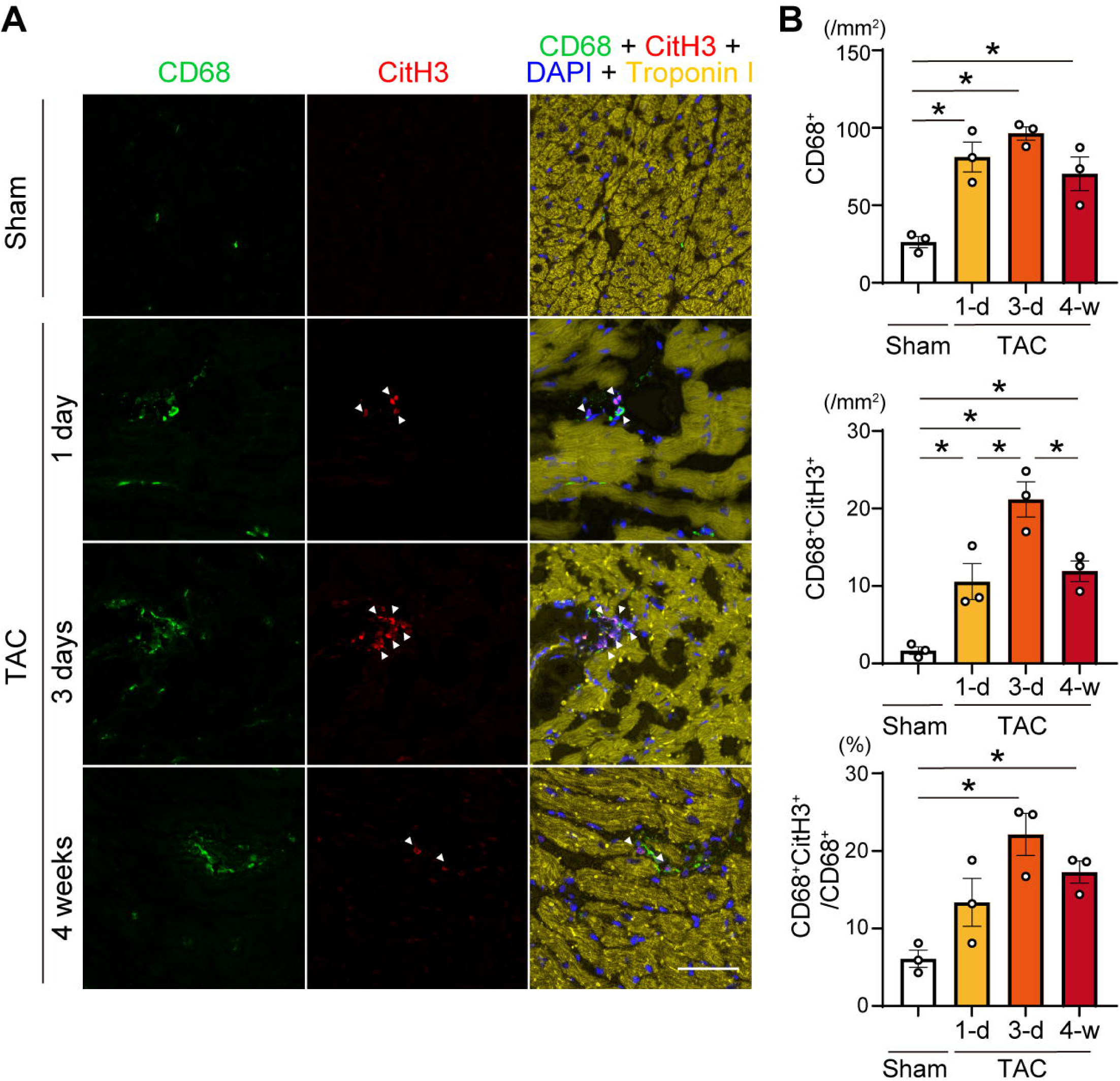
Induction of myocardial METs in response to pressure overload in mice. **A**, Left ventricular tissue sections from wild-type mice were immunostained for CD68 (green), citrullinated histone H3 (CitH3, red), troponin I (yellow) and DAPI (blue) at 1, 3 days, and 4 weeks post-transverse aortic constriction (TAC) and in sham-operated controls. Arrowheads indicate MET-positive cells (CD68^+^CitH3^+^). Scale bars, 50 μm. **B**, Quantification of CD68^+^ macrophages, METs (CD68^+^CitH3^+^) per tissue area, and MET-to-macrophage ratio (CD68^+^CitH3^+^/CD68^+^) (n=3 in each). All data are presented as mean ± SEM. *P < 0.05 by one-way analysis of variance with Tukey’s post hoc analysis.

### 3.3 PAD4 is involved in MET formation

Peptidyl arginine deiminase 4 (PAD4) plays a crucial role in extracellular trap formation by catalyzing the citrullination of arginine residues in histone proteins, thereby reducing their positive charge and promoting chromatin decondensation.^21^ To examine the contribution of PAD4 to MET formation, we investigated its role in the macrophage cell line RAW264.7. MET formation was robustly induced by several stimuli, including the calcium ionophore ionomycin, the protein kinase C activator phorbol 12-myristate 13-acetate, and lipopolysaccharide (Supplementary Figure S2). In contrast, pharmacological inhibition of PAD4 markedly suppressed MET formation (Supplementary Figure S3). We next assessed mitochondrial DNA (mtDNA) as an intrinsic trigger and found that mtDNA clearly induced METs (Figure 3A and 3B), whereas nuclear DNA failed to elicit MET formation (Supplementary Figure S4). Pharmacological inhibition of PAD4 abolished MET formation under mtDNA-stimulated conditions (Figure 3C and 3D).

**Figure 3.**
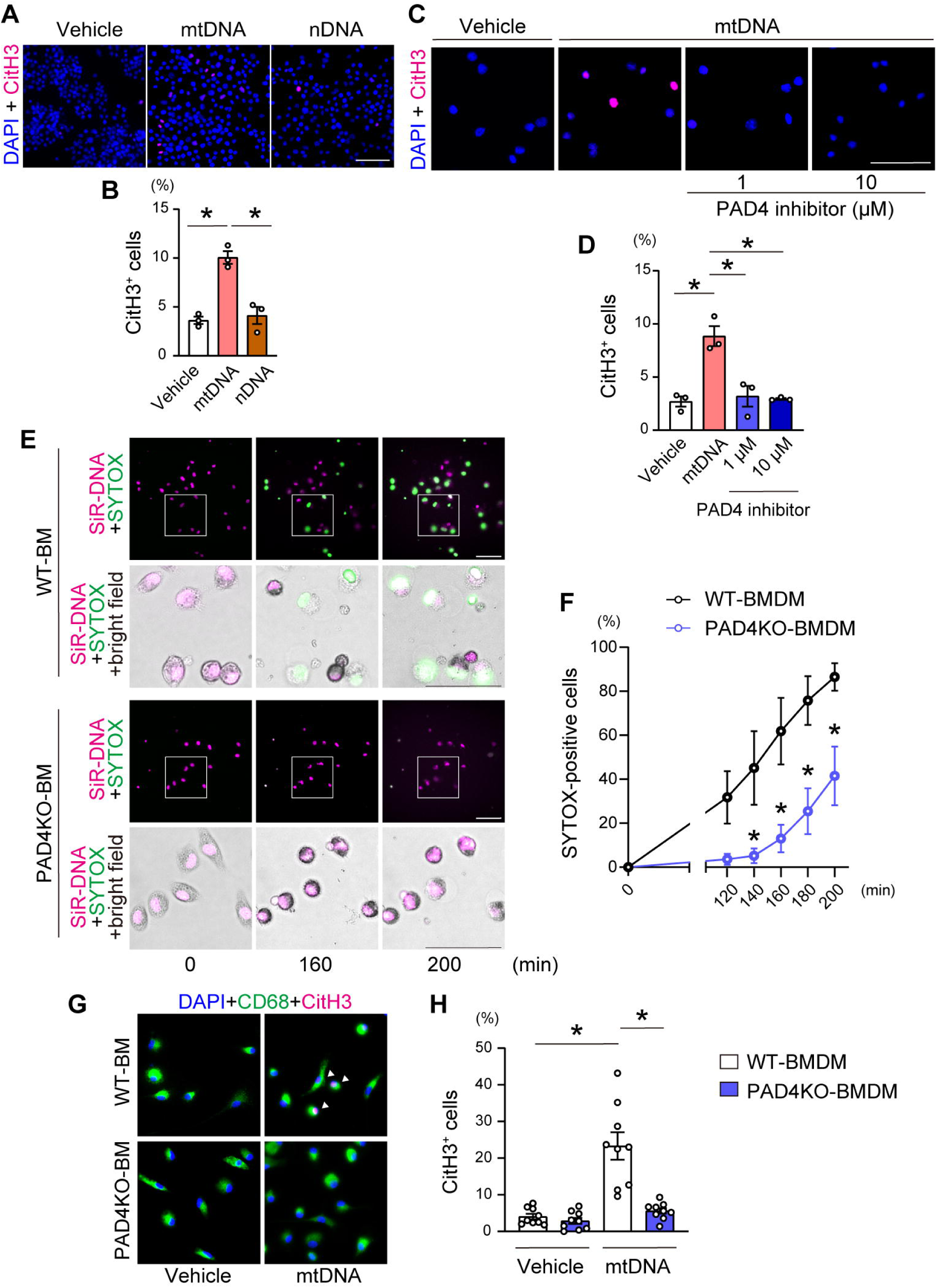
MET formation is PAD4-dependent. **A**, Representative immunofluorescence images of METs in RAW264.7 macrophage cell line. Two h after stimulation with mitochondrial DNA (mtDNA, 250 ng/mL) or nuclear DNA (nDNA, 250 ng/mL), cells were stained with an anti-citrullinated histone H3 antibody (CitH3, magenta) and DAPI (blue). Scale bars, 50 μm. **B**, Quantitative analysis of MET formation. The percentage of CitH3^+^-positive cells are shown (n=3). **C**, Representative immunofluorescence image of METs after PAD4 inhibition. Thirty min after pretreatment with the PAD4 inhibitor GSK484 at the indicated concentrations or vehicle, RAW264.7 cells were exposed to mtDNA for 2 h and stained with an anti-CitH3 antibody (magenta) and DAPI (blue). Scale bars, 50 μm. **D**, Quantitative analysis of MET formation (n=3). **E**, Time-lapse imaging of bone marrow-derived macrophages (BMDMs) from wild-type (WT) or PAD4 knockout (PAD4KO) mice following mtDNA stimulation. SiR-DNA (magenta) was used to visualize nuclei and the cell-impermeable SYTOX dye (green) was used to detect extracellular DNA. Representative images at 0, 160 and 200 min are shown. Images at higher magnification are displayed from boxed area in the lower panels. Scale bars, 50 μm. **F**, Quantification of SYTOX-positive cells (n=4). *P < 0.05 by two-way repeated-measures analysis of variance followed by Sidak’s multiple comparisons. **G**, WT or PAD4KO BMDMs were stimulated with mtDNA for 6 h, fixed, and stained with anti-CD68 (green) and anti-CitH3 (magenta) antibodies and DAPI (blue). Representative immunofluorescence images are shown. Arrowheads indicate CitH3-positive METs. **H**, Quantification of MET formation (n=3). *P < 0.05 by one-way analysis of variance with Tukey’s post hoc analysis. All data are presented as mean ± SEM.

Next, we used PAD4KO mice and confirmed reduced PAD4 protein levels in monocytes/macrophages from bone marrow cells and circulating leukocytes (Supplementary Figure S5). Time-lapse live-cell imaging demonstrated that BMDMs from WT mice developed METs in a time-dependent manner in response to mtDNA stimulation, whereas BMDMs from PAD4KO mice showed significantly reduced MET formation compared with WT BMDMs at all time points examined after 130 min (Figure 3E and 3F). Consistently, under fixed conditions, the number of MET-forming CD68^+^ BMDMs from PAD4KO mice was significantly lower than those from WT mice following mtDNA stimulation (Figure 3G and 3H). Collectively, these findings indicate that PAD4 is required for MET formation.

### 3.4 Bone marrow-derived PAD4 deficiency attenuates cardiac remodelling and improves survival in a pressure overload murine model

To determine the in vivo contribution of PAD4 to METs in heart failure, BMT was performed using bone marrow cells from PAD4KO mice (Figure 4A). Four weeks after BMT, the recipient mice were then subjected to sham or TAC surgery and followed for an additional four weeks. Echocardiographic analysis revealed that TAC significantly induced enlargement of LV diameter at both diastole and systole and reduced fractional shortening in WT-BMT mice, while PAD4KO-BMT mice displayed smaller LV diameters and preserved fractional shortening (Figure 4B and 4C). Although TAC increased diastolic interventricular septal and posterior wall thickness in both WT-BMT and PAD4KO-BMT mice, no significant differences were observed between the groups. The whole heart weight and LV weight normalized to tibial length did not differ significantly between groups (Figure 4D). In contrast, the lung weight-to-tibial length ratio was significantly lower in PAD4KO-BMT mice compared to WT-BMT mice, indicating a protective effect of PAD4 deficiency against TAC-induced pulmonary congestion. Consistent with these findings, PAD4KO-BMT mice subjected to TAC exhibited improved survival compared with WT-BMT mice (Figure 4E). Histological assessment demonstrated increased interstitial fibrosis in both WT-BMT and PAD4KO-BMT hearts following TAC, while PAD4KO-BMT mice displayed a significantly lower myocardial fibrosis fraction than WT-BMT mice (Figure 4F and 4G). The cross-sectional area of cardiomyocytes was increased in response to TAC, but did not differ between WT-BMT and PAD4KO-BMT mice (Figure 4H). Expression levels of BNP, as well as Collagen III were elevated in WT-BMT mice but those were significantly lower in PAD4KO-BMT than WT-BMT mice (Figure 4I). Together, these findings suggest that deletion of PAD4 in bone marrow cells prevents pressure overload-induced left ventricular systolic dysfunction and cardiac remodelling, thereby improving survival.

**Figure 4.**
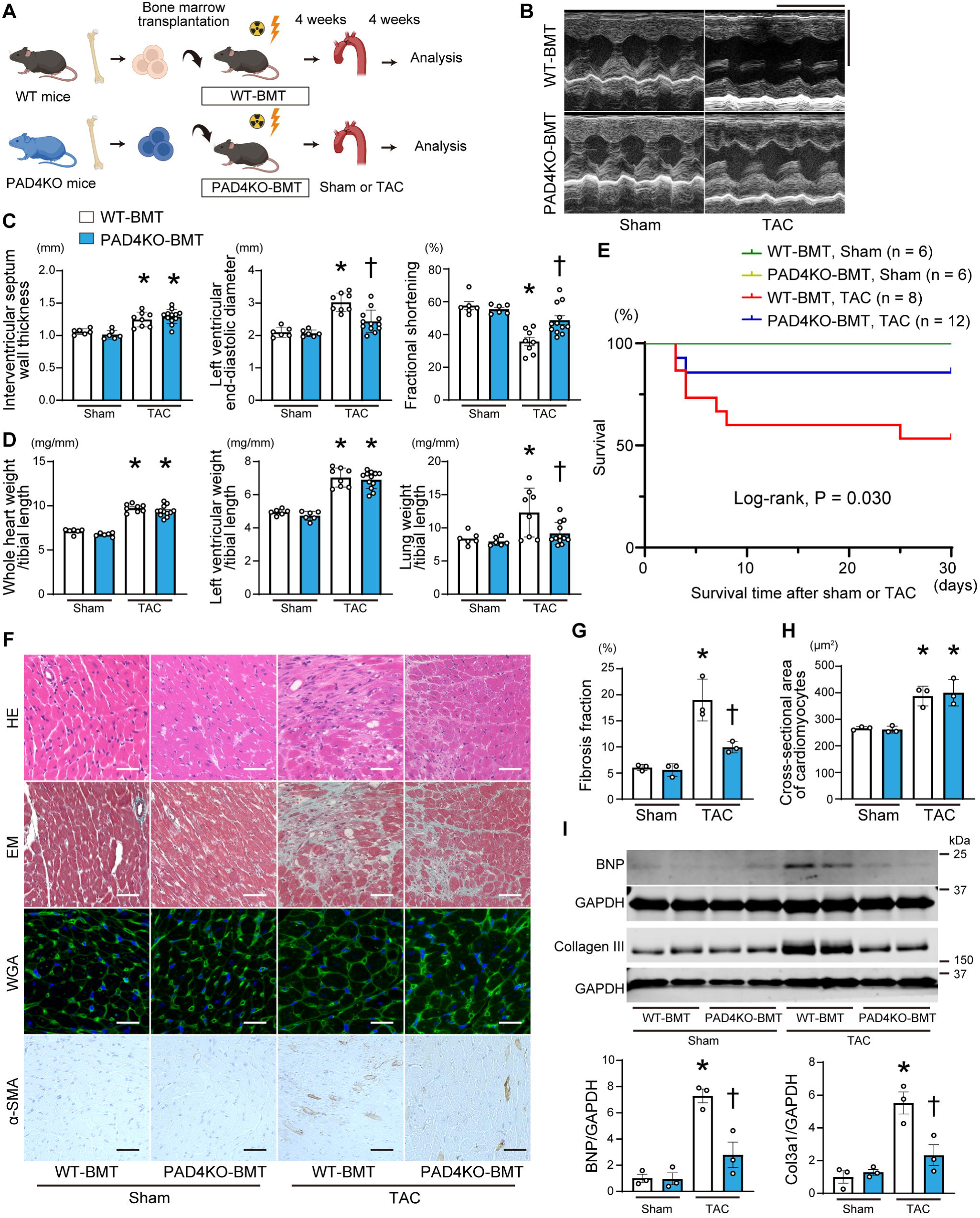
Bone marrow-derived PAD4 deficiency attenuates cardiac remodelling and improves survival in a mouse model of heart failure. **A**, Schematic representation of the experimental design. Bone marrow cells from wild-type (WT) or PAD4 knockout (KO) mice were injected into lethally irradiated mice. Four weeks after bone marrow transplantation (BMT), the recipient mice (WT-BMT or PAD4KO-BMT) underwent a sham procedure or transverse aortic constriction (TAC) and were analysed 4 weeks after the operation. **B**, Representative images of M-mode echocardiograms. Scale bars, 0.2 seconds and 4 mm, respectively. **C**, Echocardiographic assessment. Sham-operated WT-BMT mice (n=6), sham-operated PAD4KO-BMT mice (n=6), TAC-operated WT-BMT mice (n=8), and TAC-operated PAD4KO-BMT mice (n=12). **D**, Physiological parameters. **E**, Kaplan–Meier survival curves of WT-BMT and PAD4KO-BMT mice following either sham or TAC surgery. The number of animals in each group and the P value from the log-rank test are indicated. **F**, Histological analyses with representative images are shown. Heart sections from WT-BMT and PAD4KO-BMT mice were stained with Hematoxylin–Eosin (HE), Elastica–Masson (EM), and wheat germ agglutinin (WGA), or immunostained with an anti-αSMA antibody. Scale bars, 50 μm. **G**, Quantitative analysis for fibrosis fraction in EM–stained sections (n=3 in each). **H**, Quantification of the cross-sectional area of cardiomyocytes in WGA-stained sections (n=3 in each). **I**, Immunoblot analysis of BNP and Collagen III in heart extracts using the indicated antibodies. Densitometric quantification is shown in the panels, with GAPDH as the loading control. The mean value of sham-operated WT-BMT mice was normalized to 1 (n=3 in each). All data are presented as mean ± SEM. *P < 0.05 versus the corresponding sham-operated group, and ^†^P < 0.05 versus TAC-operated WT-BMT mice as determined by one-way analysis of variance with Tukey’s post hoc analysis.

### 3.5 Bone marrow-derived PAD4 deficiency prevents MET formation in myocardial tissue

Immunohistochemical analyses demonstrated a robust increase in METs in the left ventricle of WT-BMT mice at 4 weeks after TAC, whereas MET formation was markedly attenuated in PAD4KO-BMT mice (Figure 5A and 5B). Consistent with this reduction in METs, the number of CD68-positive macrophages was also significantly lower in PAD4KO-BMT than in WT-BMT hearts following TAC. To further define the early inflammatory response to pressure overload, we examined left ventricular tissue at 3 days post-TAC. At this early time point, WT-BMT hearts already exhibited substantial MET accumulation, while METs remained nearly undetectable in PAD4KO-BMT hearts (Supplementary Figure S6). Collectively, these findings indicate that PAD4 deficiency in bone marrow-derived cells suppresses cardiac MET formation during both the early and late phases of TAC-induced pressure overload.

**Figure 5.**
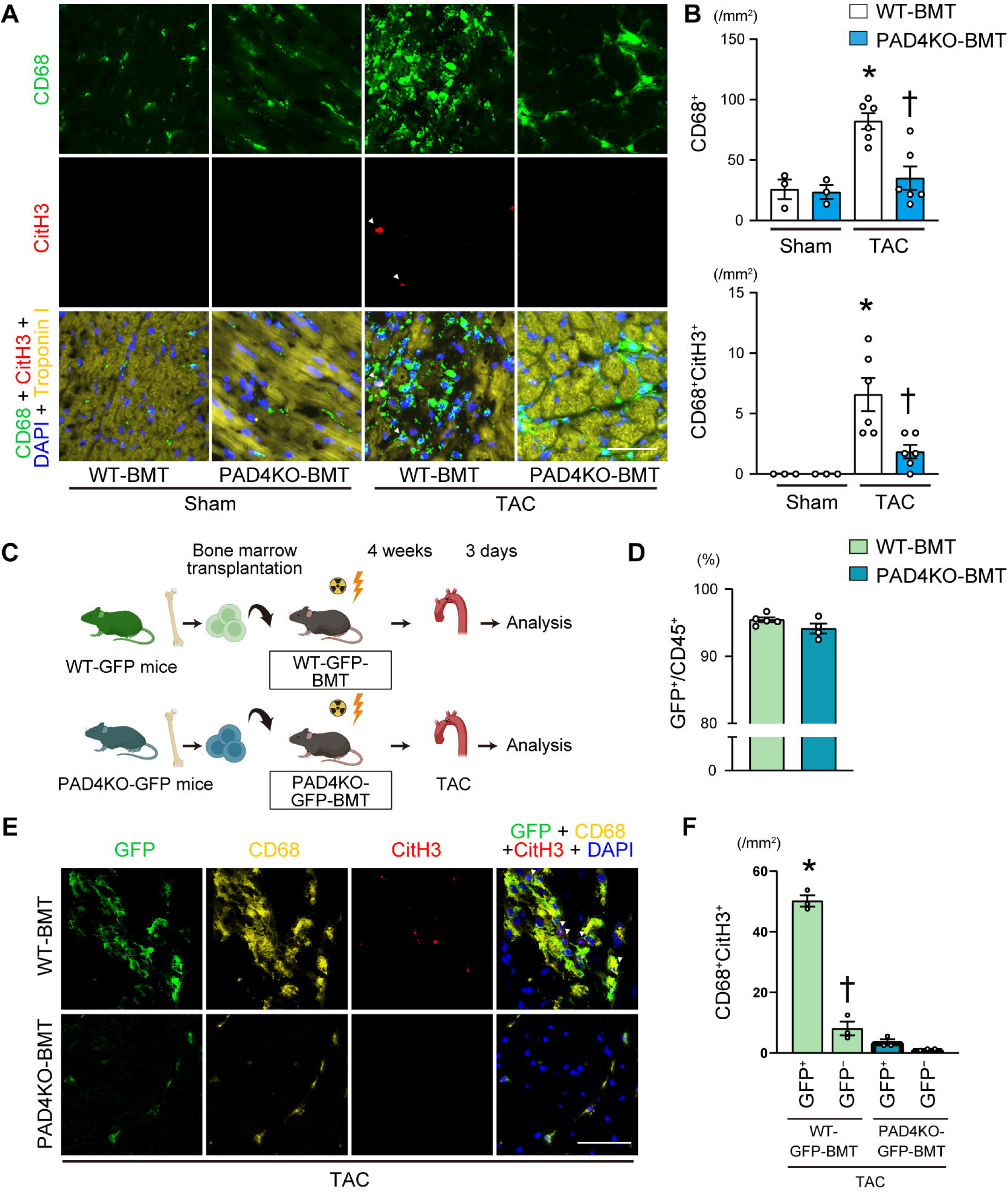
Hematopoietic PAD4 deficiency suppresses MET formation in pressure-overload hearts. **A**, Representative fluorescent immunohistochemistry images of left ventricular tissue sections from WT-BMT and PAD4KO-BMT mice 4 weeks after sham or TAC surgery. The sections were stained with antibodies against CD68 (green), citrullinated histone H3 (CitH3, red), and troponin I (yellow), along with nuclear staining using DAPI (blue). Scale bars, 50 μm. **B**, Quantification of CD68^+^ macrophages and METs (CD68^+^CitH3^+^) per tissue area (n=3-6). *P < 0.05 versus the corresponding sham-operated group, and ^†^P < 0.05 versus TAC-operated WT-BMT mice as determined by one-way analysis of variance with Tukey’s post hoc analysis. **C,** Schematic illustration of the experimental design using GFP-expressing mice. Bone marrow cells from WT-GFP or PAD4KO-GFP donor mice were transplanted into irradiated mice. The chimeric recipient mice (WT-GFP-BMT or PAD4KO-GFP-BMT) underwent TAC surgery and were analysed 3 days later. **D**, Chimerism analysis of the percentages of GFP^+^ cells within CD45^+^ cells in the peripheral blood by flow cytometry, 4 weeks after BMT. **E**, Representative immunofluorescent images of the left ventricular tissue in WT-GFP-BMT or PAD4KO-GFP-BMT mice. The sections were stained with antibodies against CD68 (yellow) and citrullinated histone H3 (CitH3, red) along with DAPI (blue). GFPL cells were detected using GFP fluorescence. Arrowheads indicate GFP^+^CD68^+^CitH3^+^ cells. Scale bars, 50 μm. **F**, Quantification of GFP-positive and GFP**-**negative CD68^+^CitH3^+^ cells. *P < 0.05 versus all other groups, and ^†^P < 0.05 versus GFP-negative CD68^+^CitH3^+^ in PAD4KO-GFP-BMT mice, as determined by one-way analysis of variance with Tukey’s post hoc analysis. All data are presented as mean ± SEM.

To distinguish the contribution of BMDMs from that of resident cardiac macrophages, we visualized bone marrow-derived cells using CAG-EGFP mice^14^ and crossed them with PAD4KO mice or WT mice. Bone marrow cells from these mice were transplanted into lethally irradiated WT mice, and four weeks later, the mice were subjected to TAC surgery (Figure 5C). FACS analysis showed the majority of hematopoietic cells in peripheral blood were GFP-positive, indicating engraftment of donor-derived bone marrow cells (Figure 5D and Supplementary Figure S7). Immunohistochemical analysis revealed that the majority of CitH3^+^CD68^+^ macrophages were GFP-positive (Figure 5E and 5F), suggesting that infiltrating BMDMs are a key source of METs. In contrast, PAD4KO-BMT hearts showed reduced CitH3-positive GFP^+^CD68^+^ macrophages. These findings suggest that bone marrow-derived PAD4 is essential for MET formation in the heart after pressure overload and is closely associated with cardiac remodelling.

### 3.6 PAD4 deficiency protects against cardiac remodelling via neutrophil-independent mechanisms

We next investigated the differential contributions of PAD4-dependent NET and MET formation to TAC-induced HFrEF. To this end, we depleted neutrophils using an anti-Ly6G-based approach (Figure 6A).^18,19^ Administration of the anti-Ly6G antibody significantly reduced peripheral white blood cell counts and the neutrophil fraction compared to IgG isotype controls in both WT-BMT and PAD4KO-BMT mice after TAC, without affecting monocyte-macrophage abundance in either group (Figure 6B and Supplementary Figure S8). Echocardiographic analysis at 4 weeks post-TAC demonstrated that IgG-injected PAD4KO-BMT mice displayed higher fractional shortening than WT-BMT mice (Figure 6C), consistent with our earlier findings. Lung weight tended to be lower in anti-Ly6G antibody-injected PAD4KO-BMT mice than in IgG-injected PAD4KO-BMT mice (Supplementary Figure S9). Anti-Ly6G treatment improved fractional shortening in WT-BMT mice compared to IgG-injected WT-BMT mice, indicating a pathogenic role for neutrophils in TAC-induced HFrEF. Importantly, PAD4KO-BMT mice receiving anti-Ly6G treatment displayed an even greater improvement in fractional shortening than anti-Ly6G-treated WT-BMT mice, indicating that PAD4 deficiency confers additional protection beyond the contribution of neutrophils and that neutrophil-independent mechanisms are involved in TAC-induced systolic dysfunction.

**Figure 6.**
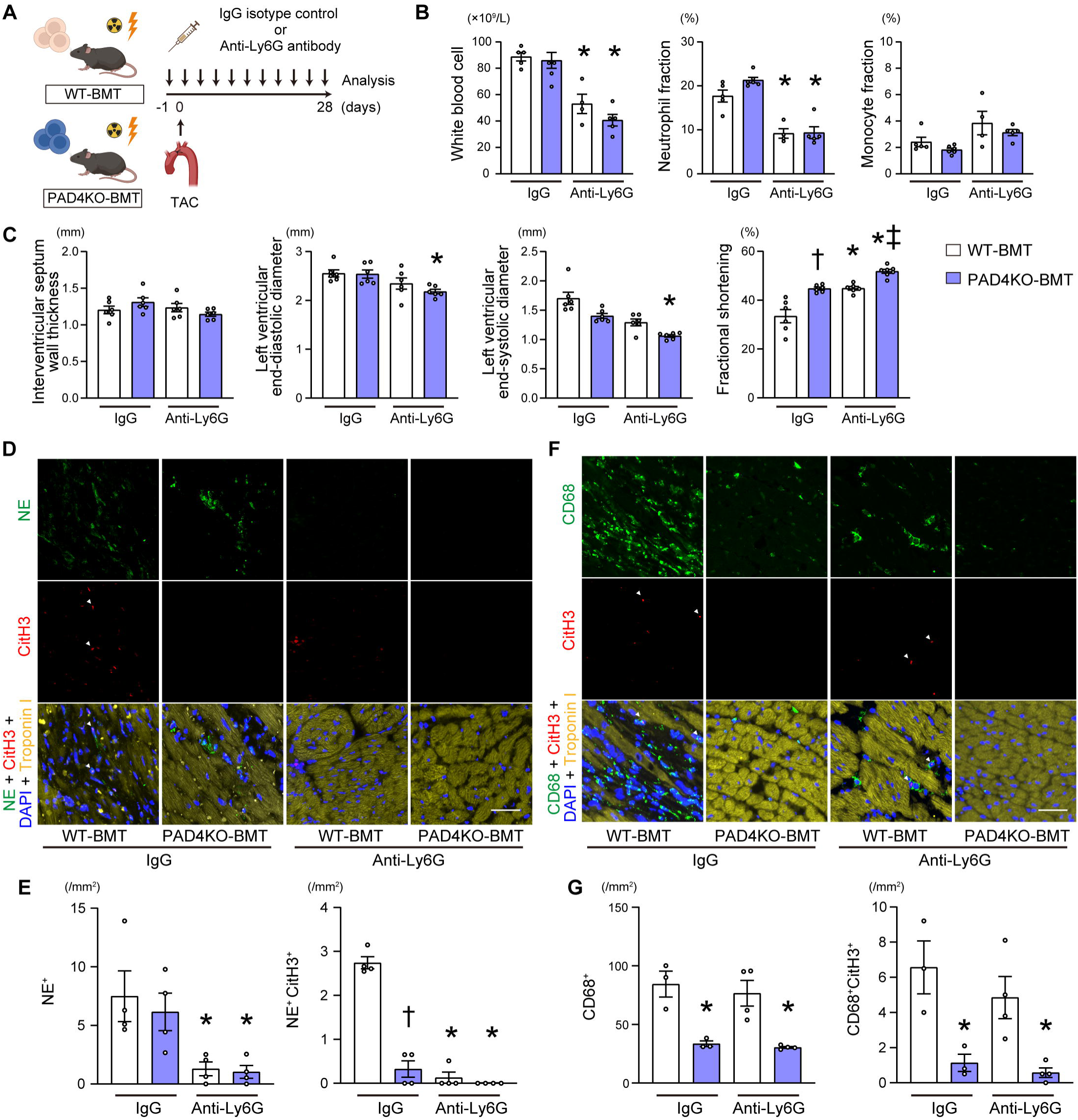
PAD4 deficiency protects against cardiac remodelling via neutrophil-independent mechanisms. **A**, Schematic overview of the experimental design. WT-BMT and PAD4KO-BMT mice received either IgG isotype control or anti-Ly6G antibody (100 μg per mouse) by intraperitoneal injection every 3 days for 28 days, as indicated. **B**, Peripheral blood cell counts, and neutrophil and monocyte fractions (n=4-5). Neutrophils were determined by CD45^high^ Gr-1^high^ CD11b^low^ and monocytes were determined by CD45^high^ Gr-1^low^ CD11b^high^ by FACS analysis. **C**, Echocardiographic assessment 28 days after TAC (n=6 in each group). **D**, Representative fluorescent immunohistochemistry images of left ventricular tissue stained for neutrophil elastase (NE, green), citrullinated histone H3 (CitH3, red), and troponin I (yellow), with nuclear counterstaining by DAPI (blue). Arrowheads indicate NET-positive cells (NE^+^CitH3^+^). Scale bars, 50 μm. **E**, Quantification of the numbers of neutrophils (NE^+^) and NETs (NE^+^CitH3^+^) per myocardial tissue area (n=4). **F**, Representative left ventricular sections stained for CD68 (green), CitH3 (red), troponin I (yellow) and DAPI (blue). Arrowheads indicate MET-positive cells (CD68^+^CitH3^+^). Scale bars, 50 μm. **G**, Quantification of CD68^+^ macrophages and METs (CD68^+^CitH3^+^) per tissue area (n=3-4). All data are presented as mean ± SEM. *P < 0.05 versus the corresponding IgG-treated group, ^†^P < 0.05 versus IgG-treated WT-BMT mice, and ^‡^P < 0.05 versus anti-Ly6G-treated WT-BMT mice as determined by one-way analysis of variance with Tukey’s post hoc analysis.

Immunohistochemical analysis revealed that the numbers of neutrophil elastase (NE)-positive neutrophils did not differ between IgG-treated WT-BMT and PAD4KO-BMT hearts; however, NE^+^ citrullinated H3^+^ were significantly reduced in IgG-treated PAD4KO-BMT mice compared to IgG-treated WT-BMT mice (Figure 6D and 6E), consistent with suppressed PAD4-dependent NET formation. Anti-Ly6G treatment reduced the numbers of NE^+^ neutrophils as well as NE^+^ citrullinated H3^+^ cells in myocardial tissue in both WT-BMT and PAD4KO-BMT mice. When examining macrophages in the myocardial tissue, PAD4KO-BMT hearts in the IgG-treated groups showed reduced CD68^+^ cells and CD68^+^ citrullinated H3^+^ MET-forming cells compared to WT-BMT mice (Figure 6F and 6G), in line with our earlier observations of reduced MET formation. In contrast, anti-Ly6G treatment did not significantly alter the numbers of CD68^+^ macrophages or CD68^+^ citrullinated H3^+^ cells in either genotype. Collectively, these findings suggest that, alongside NETs, MET formation contributes to TAC-induced cardiac dysfunction, and that PAD4 deficiency confers cardio-protection, at least in part, by attenuating both processes.

### 3.7 Mitochondrial DNA-enriched exopher is a driver of METs

To further investigate the mechanisms underlying MET formation in the myocardial tissue, we focused on cardiomyocyte-derived subcellular particles reported as exophers.^20^ We used cardiomyocyte-specific tdTomato reporter mice crossed with α-myosin heavy chain promoter-driven Cre transgenic mice.^13^ Exophers were isolated from heart homogenates using fluorescence-activated cell sorting based on tdTomato expression (Figure 7A and (Supplementary Figure S10). Morphological assessment by electron microscopy indicated that these cardiac exophers were mitochondria-rich and contained high levels of mitochondrial DNA (Figure 7B and 7C). We then conducted ex vivo analysis and found that administration of the cardiac exophers induced MET formation in BMDMs (Figure 7D). These findings suggest that mitochondrial DNA-enriched exophers act as a driver of MET formation.

**Figure 7.**
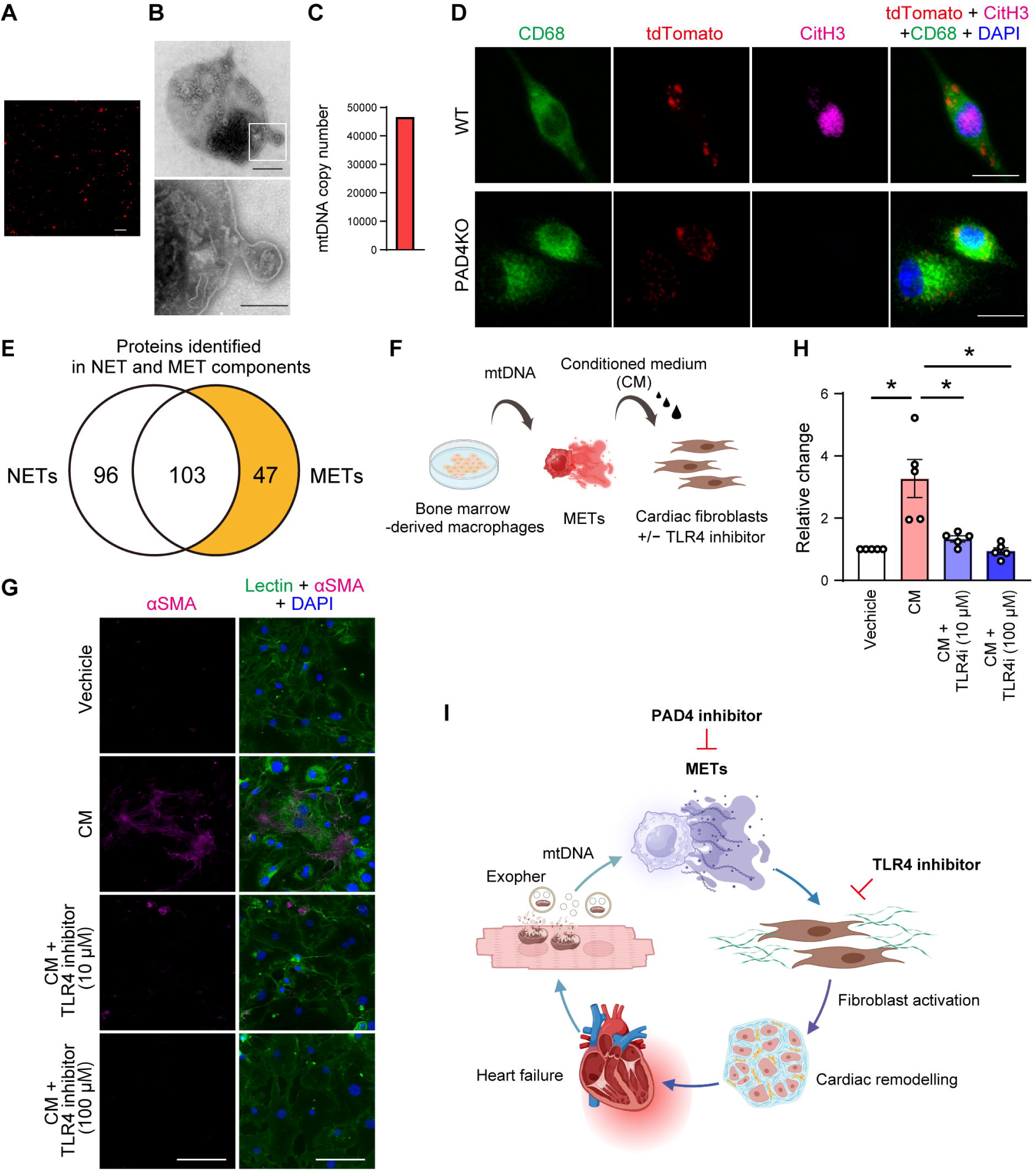
Mitochondrial DNA–enriched exophers drive METs, leading to TLR4-dependent activation of cardiac fibroblasts. **A**, Representative fluorescence images of mouse cardiomyocyte-derived exophers (tdTomato). Scale bars, 20 μm. **B**, Electron microscopy images of the exophers. The boxed area in the upper panel is enlarged and highlighted in the lower panel. Scale bars, 1 μm and 500 nm. **C**, Mitochondrial DNA (mtDNA) copy number of the cardiac exophers. **D**, Representative immunofluorescence images of BMDMs from WT or PAD4 knockout mice after exposure to cardiac exophers. At 24 h after stimulation, the cells were stained for CD68 (green), CitH3 (magenta), and DAPI (blue). Exophers were identified by endogenous tdTomato fluorescence (red). Scale bars, 10 μm. **E**, Venn diagram illustrating the number of proteins identified in NET and MET components. Peripheral neutrophils and BMDMs were stimulated with mtDNA and purified components were analysed by liquid chromatography–tandem mass spectrometry. **F**, Schematic experimental protocol. BMDMs from WT mice were stimulated with mtDNA. Primary mouse cardiac fibroblasts were incubated with the conditioned medium (CM) for 24 h in the presence or absence of a TLR4 inhibitor. **G**, Representative immunofluorescence images of cardiac fibroblasts stained with anti-αSMA antibody (magenta), wheat germ agglutinin (green) and DAPI (blue). Scale bars, 100 μm. **H**, Quantitative analysis of relative αSMA-signal intensity. Data are expressed as a relative ratio to vehicle from 5 independent experiments. All data are presented as mean ± SEM. **I**, The proposed model of METs and heart failure.

### 3.8 METs promote cardiac fibroblast activation via TLR4 pathway

Lastly, to determine how the extracellular components released during MET formation affect the resident cardiac cells, we performed proteomic profiling of the MET fraction derived from mitochondrial DNA-stimulated bone marrow macrophages. This analysis identified 150 proteins, including histones H1, H2A, H2B, H3, and H4 (Supplementary Table S5). Comparison with the NET proteome reported in our previous study^8^ revealed 47 proteins that were uniquely detected in METs (Figure 7E).

To assess the functional impact of METs, we focused on myocardial fibrosis based on our findings and stimulated primary cardiac fibroblasts with conditioned medium collected from WT BMDMs after MET induction (Figure 7F). This stimulation significantly increased α-SMA expression in fibroblasts, indicating fibroblast-to-myofibroblast transition (Figure 7G and 7H). Furthermore, pharmacological inhibition of Toll-like receptor 4 (TLR4) significantly reduced α-SMA–positive fibroblasts, indicating the TLR4-dependent fibroblast activation induced by MET-conditioned medium. Collectively, these findings suggest that PAD4-dependent MET formation from BMDMs promotes cardiac remodelling at least in part, through TLR4-mediated fibroblast activation (Figure 7I).

## 4. Discussion

The present study was the first to identify METs as a novel pathogenic driver in HFrEF pathogenesis. In patients with HFrEF, abundant METs in myocardial tissue were significantly associated with adverse LV remodelling and poor clinical outcomes. In a murine HFrEF model, bone marrow-derived hematopoietic knockout of PAD4 ameliorated cardiac remodelling and improved survival. Mechanistically, mitochondrial DNA-containing cardiomyocyte-derived exopher triggered MET formation which subsequently activated cardiac fibroblasts via TLR4 pathway. Suppressing METs represents a potential therapeutic strategy to attenuate the progression of heart failure.

Prolonged inflammation leads to impaired cardiac systolic function.^2^ Among the immune cells, macrophages play a central role in driving pathological cardiac remodelling through inflammatory responses.^22,23^ Extracellular traps have been recently recognized as key mediators of disease progression. NETs have been most extensively studied, particularly for their roles in cancer development^24^ and vascular disease biology^9,25^ including thrombosis, atherosclerosis. Other leukocyte-derived traps, eosinophil extracellular traps, have also been implicated in allergic diseases.^26^ In the hearts, we previously demonstrated the role of NETs in heart failure by inducing mitochondrial dysfunction of cardiomyocytes.^8^ In contrast, the pathological impact of METs has only recently been recognized in kidney disease^10^ and the roles of METs in the cardiovascular system had remained largely unclear prior to our present study.

Our bone marrow transplantation experiments revealed that PAD4 in hematopoietic cells, rather than in resident cells in the cardiac tissue, is required for pressure overload-induced MET formation and cardiac dysfunction. In addition, given that depletion of neutrophils partially improved cardiac dysfunction, PAD4-driven METs substantially contribute to the inflammatory and fibrotic remodelling process independent of NETs. PAD4 is unique among the PAD family because it contains a functional nuclear localization signal,^27^ and this property is central to its role in histone citrullination and extracellular trap formation. PAD4 acts as a shared upstream regulator of both neutrophil- and macrophage-derived extracellular traps, suggesting that selective inhibition of PAD4 or trap formation could provide a broad anti-inflammatory strategy for heart failure. Although citrullinated histones are pro-inflammatory, differential impacts of METs and NETs have been demonstrated by our proteomic approach. The components from NETs directly impair mitochondrial respiration in cardiomyocytes,^8^ whereas those from METs promote fibroblast activation and cardiac remodelling, creating a vicious cycle that exacerbates cardiac dysfunction and contributes to the pathogenesis of HFrEF. Citrullinated histones released by METs can trigger intracellular signalling in recipient cells^28^ by binding to and activating TLR4.^29^ Cardiac fibroblast activation is considered as a major process of myocardial fibrosis and remodelling,^30^ and immune-fibroblast cell communication has been increasingly recognized.^31^ Further investigation is required to determine the specific effects of MET contents, beyond citrullinated histones, on cardiac fibroblasts and other cell types.

Our study indicates that extracellular vesicles derived from cardiomyocytes serve as potent inducers of MET formation. Exophers are approximately 3.5 µm in diameter, larger than conventional exosomes, and are enriched in mitochondrial components including mitochondrial DNA. Given that mitochondrial DNA harbors CpG motifs similar to bacterial DNA, it can trigger TLR9 signalling.^32^ Pressure overload-induced cardiomyocytes release mitochondrial DNA-enriched exophers, which activate PAD4 in macrophages and promote MET formation, but the precise molecular mechanisms remain to be clarified.

From a translational perspective, identifying METs in the myocardium in patients with HFrEF may help predict adverse outcomes and enhance risk stratification. Therapeutic strategies targeting PAD4 and TLR4 may attenuate cardiac remodelling and progression to heart failure.

## Study limitations

The present study has several limitations. The clinical investigation was conducted at a single center with a modest sample size and a limited number of events, which may reduce the generalizability of the findings. Since only patients who underwent endomyocardial biopsy were enrolled, the possibility of selection bias and the impact of unmeasured confounding variables cannot be excluded. In the animal experiments, we employed a bone marrow transplantation model with hematopoietic cells derived from PAD4-deficient male mice; therefore, future studies using macrophage-specific PAD4 deletion models will be needed to confirm the cell-specific role of PAD4.

## Conclusions

PAD4-dependent MET formation from BMDMs represents a novel driver of cardiac remodelling. Targeting MET formation may provide a potential therapeutic strategy for heart failure.

## Authors’ contributions

SI and TM designed the research, performed experiments and analysed the data and wrote the manuscript. SO, RO, ShM, and TY contributed to the experiments and data analysis. SaM and KU performed the flow cytometry experiments and analysed the results. YuS, TS, and MO collected the clinical data and analysed patient data. AY, KI, TI, and YT contributed to the study design, revised the manuscript, and supervised the project.

## Funding

This work was supported by the Japan Society for the Promotion of Science KAKENHI grants 23K07556 to SI and 23K27600 to YT, and SENSHIN Medical Research Foundation to TM.

## Acknowledgements

The authors gratefully acknowledge Tomiko Miura, Tomomi Ogata, Kumiko Watanabe, Yumi Yoshihisa, and Shiori Togashi for their technical assistance; Toshiyuki Suzuki for his exceptional assistance with liquid chromatography–tandem mass spectrometry analysis; and Hiiro Sannohe for valuable technical support. Figures were created with BioRender.com.

## Conflict of Interest

TY and TS belong to a department supported by Minamisoma City. The remaining authors have no disclosures to report.

## Data availability

The data for this study are available in the article and its online supplementary materials.

