## Supplemental Material for "Macrophage extracellular traps promote maladaptive cardiac remodelling and heart failure via PAD4-dependent mechanisms"

### - Figure S1-S10

### - Table S1-S5

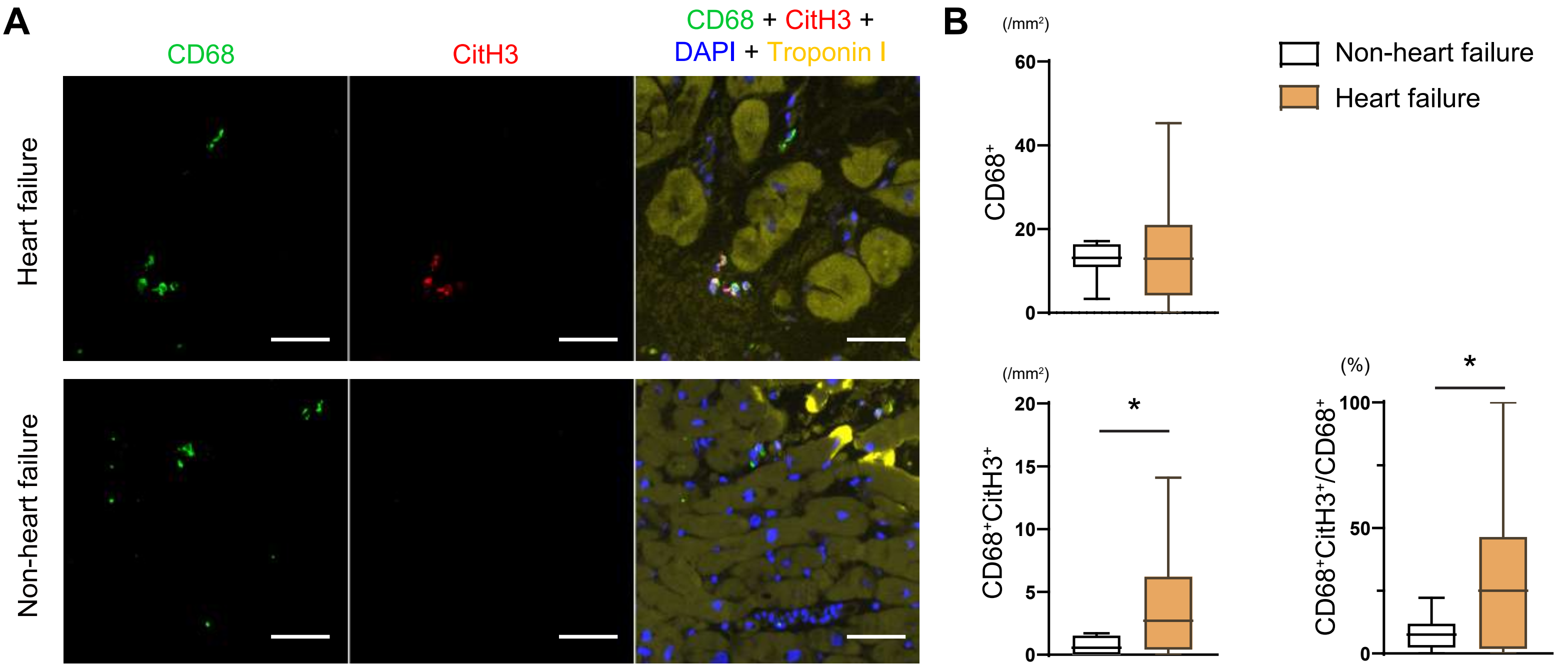


**Supplementary Figure S1. Macrophage extracellular traps (METs) on human myocardial biopsy specimens in patients with heart failure and non-heart failure.**

A, Representative fluorescent immunohistochemistry images of myocardial biopsy specimens showing the identification of METs using antibodies against CD68 (green), citrullinated histone H3 (CitH3, red), and troponin I (yellow), along with nuclear staining using DAPI (blue). Images from the heart failure group are the same as those shown in Figure 1A. Scale bars, 50 μm B, Quantification of the numbers of $CD68^+$ macrophages and METs ($CD68^+CitH3^+$) per myocardial tissue area, and MET-to-macrophage ratio ($CD68^+CitH3^+/CD68^+$). *P < 0.05 by Mann-Whitney U test.

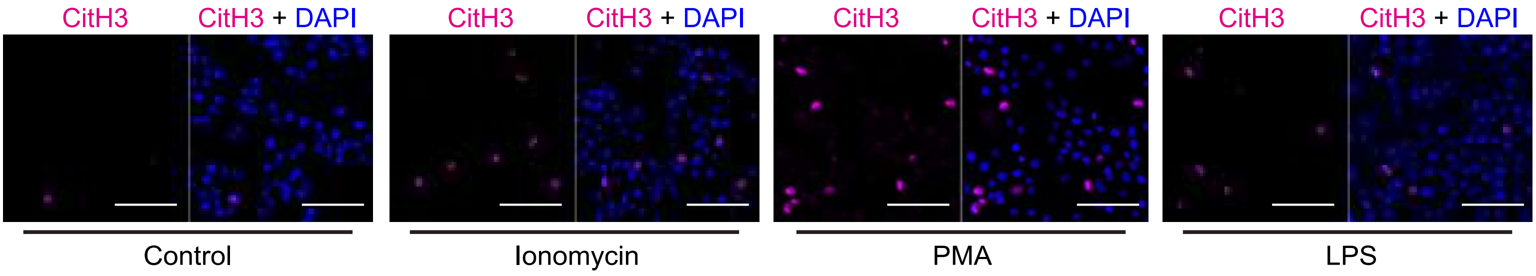


**Supplementary Figure S2. Induction of METs in RAW264.7 cells.**

Representative immunofluorescence images of METs in RAW264.7 macrophage cell lines. Cells were stimulated with ionomycin (5 μM), phorbol 12-myristate 13-acetate (PMA, 50 μM), or lipopolysaccharide (LPS, 100 ng/mL) for 6 h, then fixed and stained with an anti–citrullinated histone H3 (CitH3, magenta) antibody and DAPI (blue). Scale bars, 50 μm. Representative images from two independent experiments are shown.

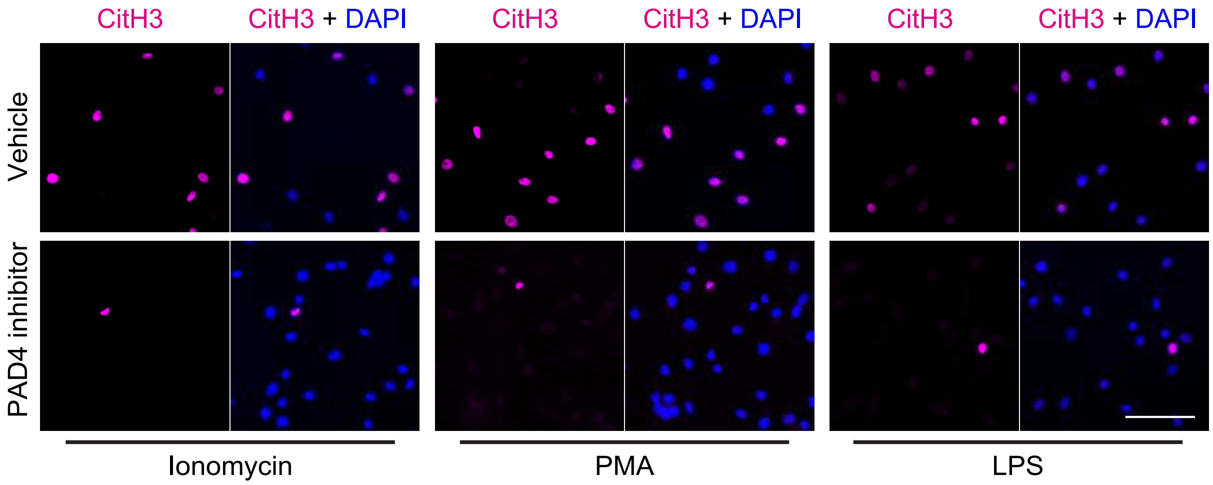


**Supplementary Figure S3. Pharmacological inhibition of PAD4 reduces the induction of METs.**
Representative immunofluorescence image of METs in RAW264.7 macrophages. Cells were pretreated with the PAD4 inhibitor GSK484 (10 μM) or vehicle for 30 min and then stimulated with ionomycin (5 μM), phorbol 12-myristate 13-acetate (PMA, 50 μM), or lipopolysaccharide (LPS, 100 ng/mL) for 6 h. Cells were fixed and stained with an anti–citrullinated histone H3 (CitH3, magenta) antibody and DAPI (blue). Scale bars, 50 μm. Representative images of two independent experiments are shown.

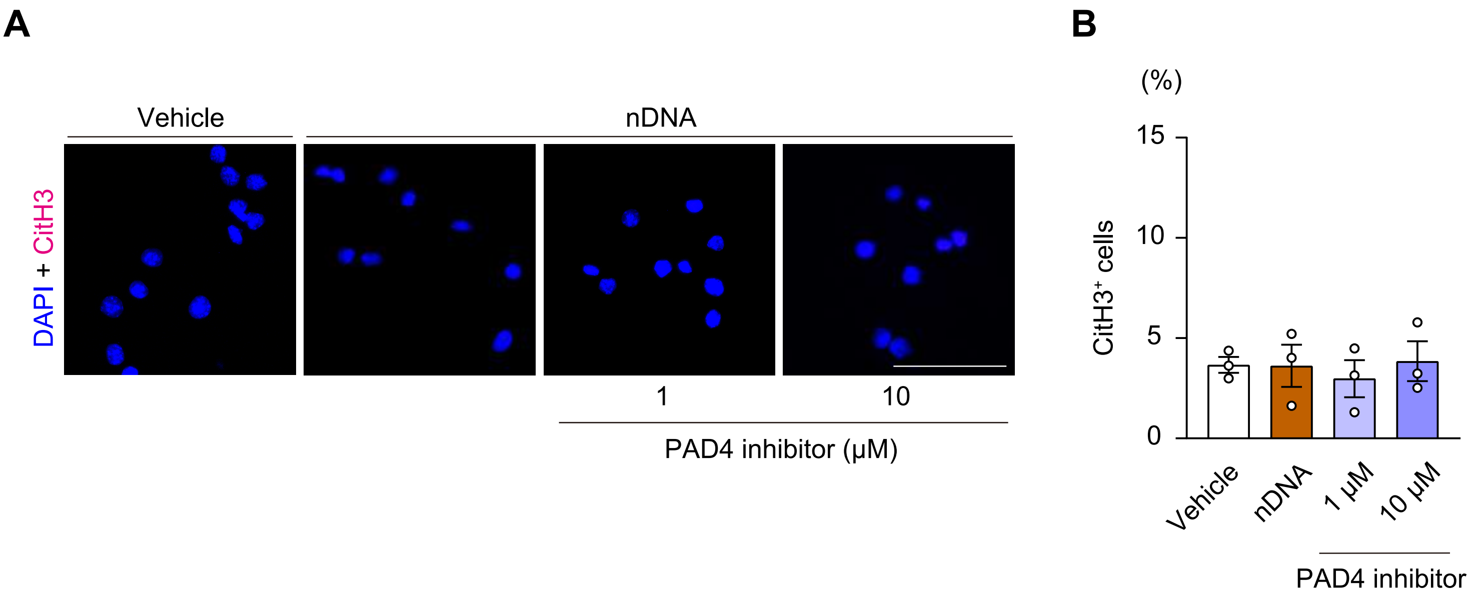


**Supplementary Figure S4. Nuclear DNA stimulation has no effect on MET formation.**
**A**, Representative immunofluorescence image showing that nuclear DNA stimulation does not induce MET formation in RAW264.7 cells, with or without PAD4 inhibition. Thirty min after pretreatment with the PAD4 inhibitor GSK484 at the indicated concentrations or vehicle, RAW 264.7 cells were exposed to nuclear DNA (nDNA, 250 ng/mL) for 2 h and stained with an anti-CitH3 antibody (magenta) and DAPI (blue). Scale bars, 50 μm. **B**, Quantitative analysis of MET formation (n=3) with one-way analysis of variance (p=ns).

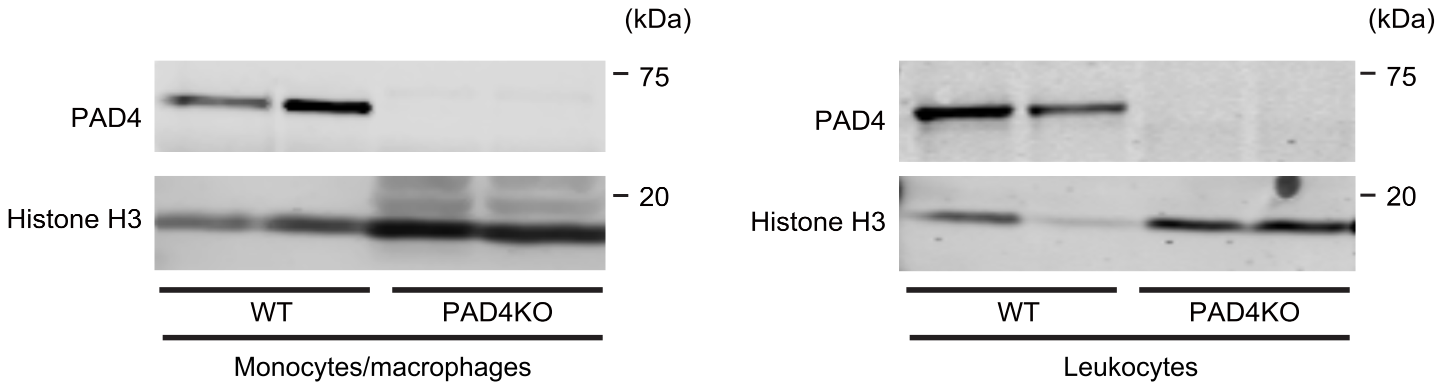


**Supplementary Figure S5. Absence of PAD4 expression in monocytes/macrophages from PAD4 knockout (KO) mice.**

Immunoblot analysis of PAD4 expression in monocytes/macrophages isolated from bone marrow cells (left panel) and peripheral blood leukocytes (right panel) of WT and PAD4KO mice. Magnetic-activated cell sorting using CD11b MicroBeads was performed to isolate myeloid cells. Histone H3 was used as a loading control. Representative images from two independent experiments are shown.

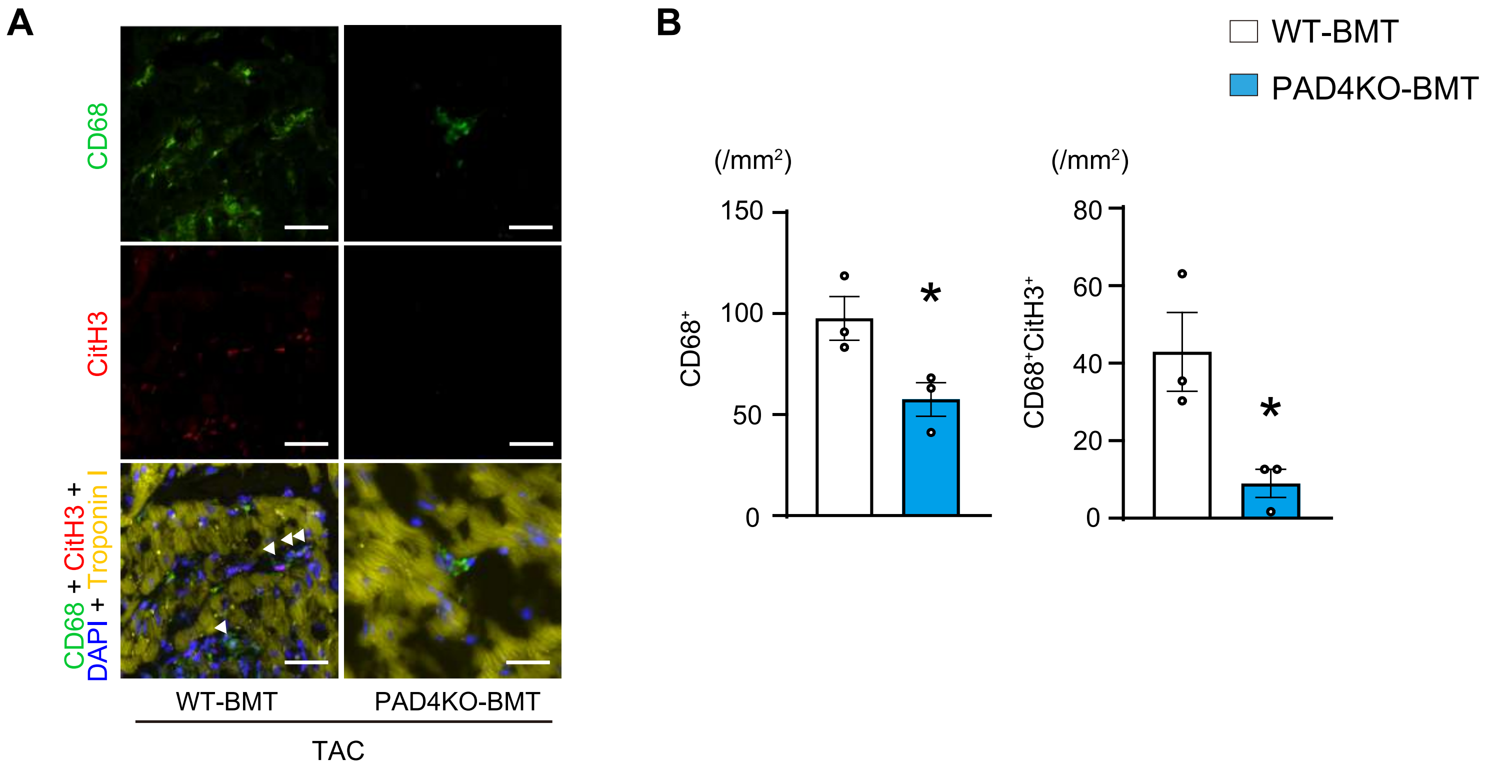


**Supplementary Figure S6. Hematopoietic PAD4 deficiency suppresses MET formation in pressure-overload hearts 3 days after transverse aortic constriction (TAC).**

**A**, Representative fluorescent immunohistochemical images of left ventricular tissue 3 days after TAC in WT-BMT and PAD4KO-BMT mice. The sections were stained with antibodies against CD68 (green), citrullinated histone H3 (CitH3, red), and troponin I (yellow), along with nuclear staining using DAPI (blue). Arrowheads indicate MET-positive cells ($CD68^{+}CitH3^{+}$). Scale bars, 50 μm. **B**, Quantification of $CD68^{+}$ macrophages and METs ($CD68^{+}CitH3^{+}$) per tissue area (n=3). *$P < 0.05$ versus WT-BMT by the unpaired t test (two-sided). All data are presented as mean ± SEM. WT-BMT indicates recipient mice transplanted with WT bone marrow cells; PAD4KO-BMT, recipient mice transplanted with PAD4 knockout bone marrow cells.

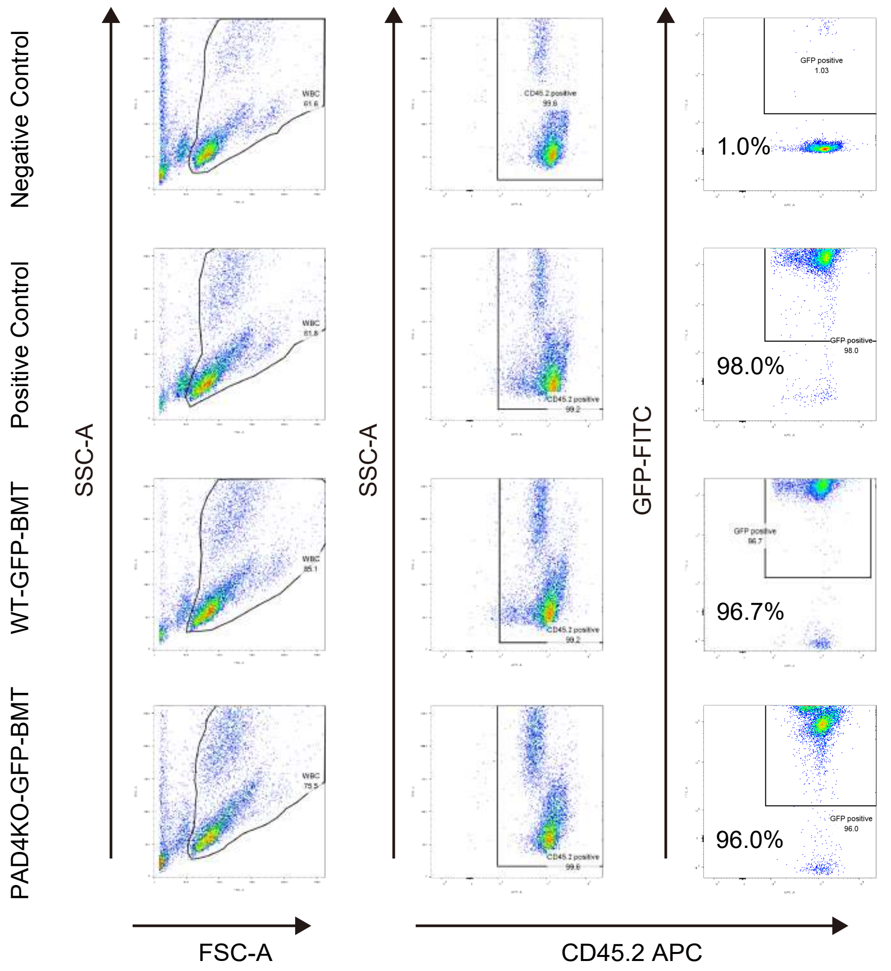


**Supplementary Figure S7. Gating strategy using GFP donor mice.**

Gating strategy for assessing peripheral blood chimerism 4 weeks after bone marrow transplantation from WT-GFP donors (WT-GFP-BMT) or PAD4-GFP donors (PAD4KO-GFP-BMT). The percentage of $GFP^{+}$ cells among circulating $CD45.2^{+}$ cells is shown in Figure 5D. WT mice were used as the negative control, and GFP mice were used as the positive control.

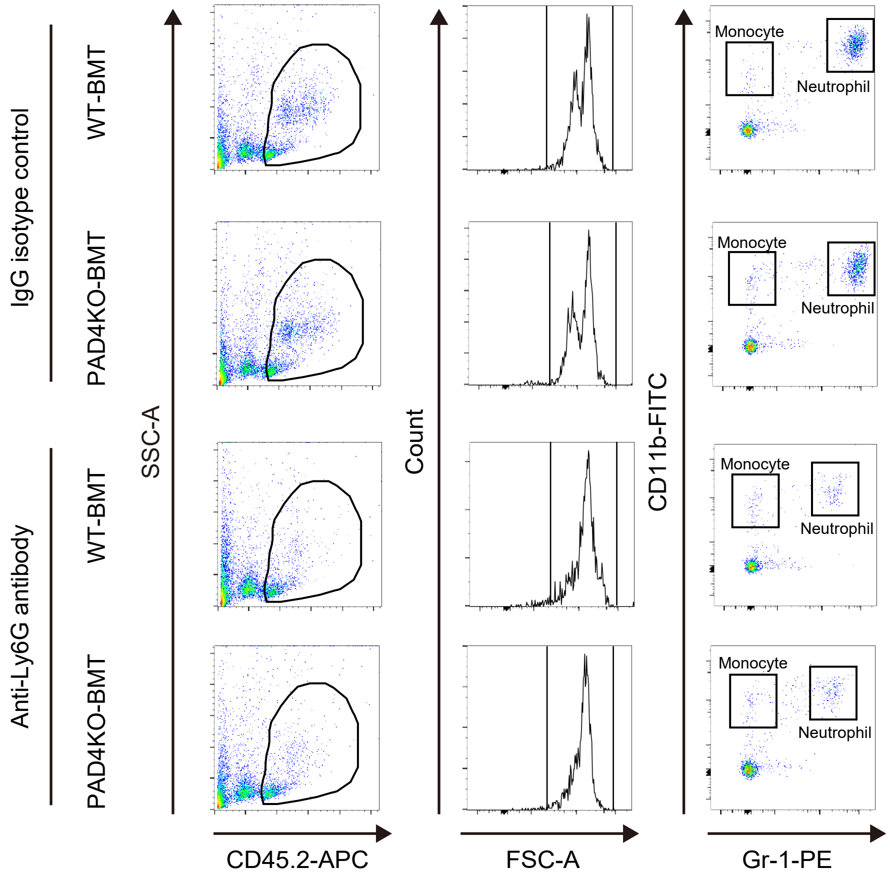


**Supplementary Figure S8. Gating strategy after an anti-Ly6G antibody administration.**

Gating strategy for analyzing neutrophil and monocyte fractions in peripheral blood after treatment with an IgG isotype control or an anti-Ly6G antibody in WT-BMT mice and PAD4KO-BMT mice. The percentages of Gr-1$^{high}$CD11b$^{high}$ (neutrophils) and Gr-1$^{high}$CD11b$^{low}$ (monocytes) among circulating CD45.2$^{+}$ cells are shown in Figure 6B.

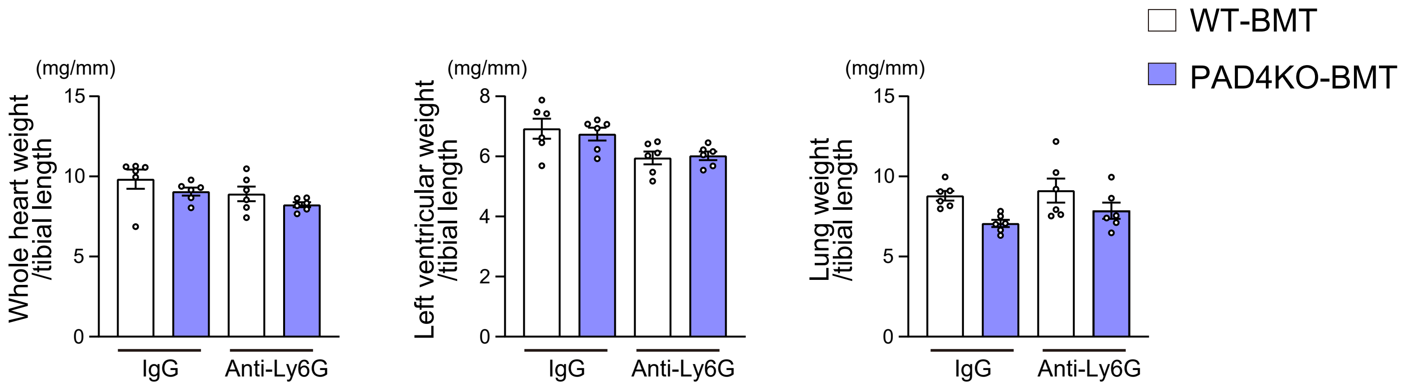


**Supplementary Figure S9. Physiological parameters.**

Whole heart weight/tibial length, left ventricular weight/tibial length, and lung weight/tibial length in WT-BMT and PAD4KO-BMT mice treated with IgG isotype control or anti-Ly6G antibody (n=6 in each group). One-way analysis of variance was used for statistical comparison, not significant.

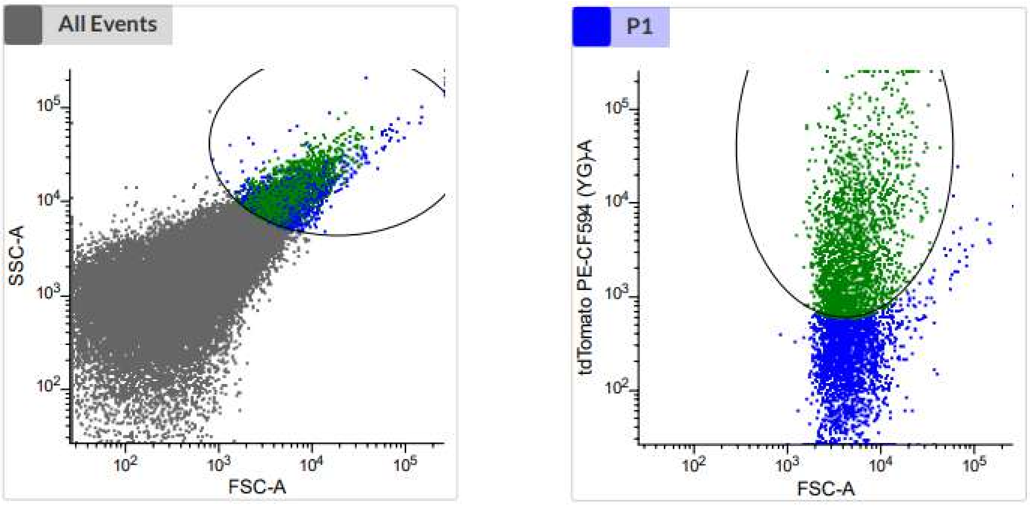


**Supplementary Figure S10. Gating strategy to isolate tdTomato-expressing cardiomyocyte-derived exophers.**
Gating strategy for cardiomyocyte-derived exophers (tdTomato).

**Table S1. Baseline characteristics of patients with heart failure with dilated cardiomyopathy (n=69).**

| | |
|---|---|
| Age, years | 55.5 ± 13.0 |
| Male sex, n (%) | 58 (84.1) |
| Body mass index, kg/$m^2$ | 24.9 ± 4.7 |
| Systolic blood pressure, mmHg | 113.2 ± 17.9 |
| Diastolic blood pressure, mmHg | 74.2 ± 11.1 |
| Smoking, n (%) | 48 (69.6) |
| NYHA functional class III/IV, n (%) | 24 (34.8) |
| **Comorbidities** | |
| Hypertension, n (%) | 31 (44.9) |
| Diabetes mellitus, n (%) | 23 (33.3) |
| Dyslipidemia, n (%) | 32 (46.4) |
| Atrial fibrillation, n (%) | 24 (34.8) |
| **Medications** | |
| Beta blockers, n (%) | 67 (97.1) |
| ACE inhibitor or ARB, n (%) | 62 (89.9) |
| Aldosterone antagonists, n (%) | 40 (58.0) |
| Diuretics, n (%) | 45 (65.2) |
| Inotropes, n (%) | 19 (30.6) |
| **Laboratory data** | |
| White blood cell, $\times 10^9$/L | 6.6 ± 1.8 |
| Absolute monocyte count, $\times 10^6$/L | 522 ± 205 |
| Hemoglobin, g/dL | 14.6 ± 1.9 |
| Albumin, g/dL | 4. 0 ± 0.6 |
| eGFR, mL/min/1.73$m^2$ | 60.7 ± 15.4 |
| Sodium, mEq/L | 140.3 ± 3.6 |
| B-type natriuretic peptide, pg/mL | 187.8 (89.2-379.1) |
| C-reactive protein, mg/dL | 0.13 (0.07-0.74) |
| **Echocardiographic data** | |
| Left ventricular diastolic diameter, mm | 62.0 ± 7.4 |
| Left ventricular systolic diameter, mm | 54.0 ± 8.2 |
| Left ventricular end-diastolic volume, mL | 199.6 ± 82.5 |
| Left ventricular end-systolic volume, mL | 148.2 ± 72.0 |
| Left ventricular ejection fraction, % | 26.9 ± 8.34 |
| Left atrial diameter, mm | 45.6 ± 7.88 |

NYHA, New York Heart Association; ACE, angiotensin-converting enzyme; ARB, angiotensin II receptor blockers; eGFR, estimated glomerular filtration rate.

**Table S2. Correlation between the numbers of METs in the myocardium and other variables.**

| | R | P value |
|---|---|---|
| Age, years | -0.182 | 0.134 |
| Body mass index, kg/m$^2$ | -0.142 | 0.198 |
| **Laboratory data** | | |
| White blood cell, ×10$^9$/L | 0.330 | 0.006 |
| Absolute monocyte count, ×10$^6$/L | 0.222 | 0.067 |
| Hemoglobin, g/dL | 0.126 | 0.302 |
| Albumin, g/dL | 0.033 | 0.799 |
| eGFR, mL/min/1.73m$^2$ | -0.237 | 0.049 |
| Sodium, mEq/L | -0.086 | 0.484 |
| B-type natriuretic peptide, pg/mL | 0.071 | 0.560 |
| C-reactive protein, mg/dL | 0.224 | 0.066 |
| **Echocardiographic data** | | |
| Left ventricular diastolic diameter, mm | 0.297 | 0.013 |
| Left ventricular systolic diameter, mm | 0.371 | 0.002 |
| Left ventricular end-diastolic volume, mL | 0.378 | 0.002 |
| Left ventricular end-systolic volume, mL | 0.374 | 0.002 |
| Left ventricular ejection fraction, % | -0.251 | 0.039 |
| Left atrial diameter, mm | 0.145 | 0.233 |
| **CPX parameters (n=43)** | | |
| Anaerobic threshold, mL/min/kg | -0.142 | 0.353 |
| Peak oxygen consumption, mL/min/kg | 0.179 | 0.212 |
| VE/$VCO_2$ slope | -0.085 | 0.558 |
| **Swan-Ganz catheterization** | | |
| PAWP, mmHg | 0.018 | 0.884 |
| Mean PAP, mmHg | 0.016 | 0.897 |
| Cardiac index, L/min/m$^2$ | 0.066 | 0.596 |
| Stroke volume index, mL/beat/m$^2$ | 0.100 | 0.420 |

METs, macrophage extracellular traps; eGFR, estimated glomerular filtration rate; CPX, cardiopulmonary exercise test; VE/$VCO_2$, minute ventilation/carbon dioxide production; PAWP, pulmonary artery wedge pressure; PAP, pulmonary artery pressure. Statistical analysis was performed by Spearman's correlation analysis.

**Table S3. Characteristics of patients stratified by the number of METs based on the optimal cutoff determined from the ROC curve.**

| | Low METs (< 2.69 /mm$^2$, n = 38) | High METs (≥ 2.69 /mm$^2$, n = 31) | P value |
|---|---|---|---|
| Age, years | 57.2 ± 12.7 | 53.5 ± 13.2 | 0.23 |
| Male sex, n (%) | 33 (86.8) | 25 (80.6) | 0.35 |
| Body mass index, kg/m$^2$ | 24.4 ± 4.9 | 25.5 ± 4.6 | 0.36 |
| Systolic blood pressure, mmHg | 116.0 ± 18.5 | 109.9 ± 16.9 | 0.16 |
| Diastolic blood pressure, mmHg | 75.4 ± 11.4 | 72.9 ± 10.7 | 0.35 |
| Smoking, n (%) | 28 (73.7) | 20 (64.5) | 0.41 |
| NYHA functional class III/IV, n (%) | 10 (26.3) | 14 (45.2) | 0.10 |
| **Comorbidities** | | | |
| Hypertension, n (%) | 20 (52.6) | 11 (35.5) | 0.15 |
| Diabetes mellitus, n (%) | 13 (34.2) | 10 (32.3) | 0.86 |
| Dyslipidemia, n (%) | 16 (42.1) | 16 (51.6) | 0.43 |
| Atrial fibrillation, n (%) | 16 (42.1) | 8 (25.8) | 0.15 |
| **Medications** | | | |
| Beta blockers, n (%) | 38 (100) | 29 (93.5) | 0.19 |
| ACE inhibitor or ARB, n (%) | 34 (89.5) | 28 (90.3) | 0.61 |
| Aldosterone antagonists, n (%) | 19 (50.0) | 21 (67.7) | 0.13 |
| Diuretics, n (%) | 25 (65.8) | 20 (64.5) | 0.91 |
| Inotropes, n (%) | 9 (23.7) | 10 (32.3) | 0.42 |
| **Laboratory data** | | | |
| White blood cell, ×10$^9$/L | 6.1 ± 1.5 | 7.2 ± 1.9 | <0.01 |
| Absolute monocyte count, ×10$^6$/L | 467 ± 161 | 590 ± 233 | 0.01 |
| Hemoglobin, g/dL | 14.3 ± 1.7 | 15.0 ± 2.1 | 0.15 |
| Albumin, g/dL | 4.0 ± 0.6 | 4.0 ± 0.6 | 0.88 |
| eGFR, mL/min/1.73m$^2$ | 63.6 ± 15.2 | 57.0 ± 15.0 | 0.07 |
| Sodium, mEq/L | 140.7 ± 3.0 | 139.7 ± 3.6 | 0.18 |
| B-type natriuretic peptide, pg/mL | 182.3 (90.3-358.9) | 187.8 (114.3-415.6) | 0.75 |
| C-reactive protein, mg/dL | 0.10 (0.04-0.34) | 0.16 (0.12-0.86) | 0.07 |
| **Echocardiographic data** | | | |
| Left ventricular diastolic diameter, mm | 60.6 ± 5.7 | 63.7 ± 8.8 | 0.08 |
| Left ventricular systolic diameter, mm | 51.9 ± 6.9 | 56.6 ± 8.9 | 0.01 |
| Left ventricular end-diastolic volume, mL | 180.3 ± 66.4 | 222.6 ± 94.3 | 0.03 |
| Left ventricular end-systolic volume, mL | 129.4 ± 51.5 | 170.7 ± 86.3 | 0.01 |

| | | | |
|---|---|---|---|
| Left ventricular ejection fraction, % | 28.5 ± 8.1 | 24.9 ± 8.3 | 0.07 |
| Left atrial diameter, mm | 45.2 ± 8.5 | 46.1 ± 7.1 | 0.65 |
| **CPX parameters** | | | |
| (low METs, n = 22; high METs, n = 21) | | | |
| Anaerobic threshold, mL/min/kg | 11.6 ± 2.4 | 11.2 ± 1.3 | 0.43 |
| Peak oxygen consumption, mL/min/kg | 16.0 ± 4.0 | 16.8 ± 3.4 | 0.42 |
| $VE/VCO_2$ slope | 33.0 ± 6.2 | 32.1 ± 5.0 | 0.57 |
| **Cardiac MRI** | | | |
| LGE-positive, n (%) | 7 (25.9) | 12 (52.2) | 0.05 |
| **Swan-Ganz catheterization** | | | |
| PAWP, mmHg | 18.3 ± 9.1 | 17.2 ± 9.6 | 0.60 |
| Mean PAP, mmHg | 25.2 ± 9.9 | 24.9 ± 12.4 | 0.92 |
| Cardiac index, L/min/$m^2$ | 2.3 ± 0.6 | 2.3 ± 0.5 | 0.95 |
| Stroke volume index, mL/beat/$m^2$ | 32.0 ± 14.7 | 33.7 ± 10.3 | 0.58 |
| **Histological analysis** | | | |
| $CD68^+$ macrophage, /$mm^2$ | 5.78 (2.13-12.15) | 15.46 (9.00-27.54) | < 0.01 |
| Ratio of METs to $CD68^+$ macrophage, % | 5.3 (0.0-25.0) | 40.0 (29.4-54.2) | < 0.01 |

NYHA, New York Heart Association; ACE, angiotensin-converting enzyme; ARB, angiotensin II receptor blockers; eGFR, estimated glomerular filtration rate; CPX, cardiopulmonary exercise test; $VE/VCO_2$, minute ventilation/carbon dioxide production; MRI, magnetic resonance imaging; LGE, late gadolinium enhancement; PAWP, pulmonary artery wedge pressure; PAP, pulmonary artery pressure; METs, macrophage extracellular traps.

**Table S4. Univariable and multivariable Cox proportional hazard analyses for cardiac events in patients with heart failure with dilated cardiomyopathy.**

| | Hazard ratio | 95% confidence interval | P value |
|---|---|---|---|
| **METs as a categorical variable** | | | |
| High METs (vs. low METs), unadjusted | 4.79 | 1.55-14.71 | 0.006 |
| High METs (vs. low METs), model 1 | 5.53 | 1.74-17.56 | 0.004 |
| High METs (vs. low METs), model 2 | 7.62 | 2.17-26.76 | 0.002 |
| **METs as a continuous variable** | | | |
| METs per 1/mm$^2$ increase, unadjusted | 1.07 | 1.00-1.15 | 0.045 |
| METs per 1/mm$^2$ increase, model 1 | 1.07 | 1.00-1.15 | 0.048 |
| METs per 1/mm$^2$ increase, model 2 | 1.10 | 1.02-1.19 | 0.014 |

Model 1: adjusted for age and sex.

Model 2: adjusted for age, sex, body mass index, and left ventricular ejection fraction.

**Table S5. Identification of protein components derived from METs by liquid chromatography–tandem mass spectrometry.**

| Number | Identified Proteins | Accession Number | Quantity |
|---|---|---|---|
| 1 | Actin, cytoplasmic 2 OS=Mus musculus GN=Actg1 PE=1 SV=1 | sp\|P63260\|ACTG_MOUSE | 2.74E+08 |
| 2 | Carbonic anhydrase 2 OS=Mus musculus GN=Ca2 PE=1 SV=4 | sp\|P00920\|CAH2_MOUSE | 1.34E+08 |
| 3 | Prothymosin alpha OS=Mus musculus GN=Ptma PE=1 SV=2 | sp\|P26350\|PTMA_MOUSE | 1.30E+08 |
| 4 | Carbonic anhydrase 1 OS=Mus musculus GN=Ca1 PE=1 SV=4 | sp\|P13634\|CAH1_MOUSE | 1.06E+08 |
| 5 | Elongation factor 1-alpha 1 OS=Mus musculus GN=Eef1a1 PE=1 SV=3 | sp\|P10126\|EF1A1_MOUSE (+2) | 8.86E+07 |
| 6 | Flavin reductase (NADPH) OS=Mus musculus GN=Blvrb PE=1 SV=3 | sp\|Q923D2\|BLVRB_MOUSE | 6.18E+07 |
| 7 | Protein S100-A9 OS=Mus musculus GN=S100a9 PE=1 SV=3 | sp\|P31725\|S10A9_MOUSE | 5.82E+07 |
| 8 | Ferritin light chain 1 OS=Mus musculus GN=Ftl1 PE=1 SV=2 | sp\|P29391\|FRIL1_MOUSE (+2) | 5.31E+07 |
| 9 | Heat shock cognate 71 kDa protein OS=Mus musculus GN=Hspa8 PE=1 SV=1 | sp\|P63017\|HSP7C_MOUSE (+5) | 5.16E+07 |
| 10 | Chitinase-like protein 3 OS=Mus musculus OX=10090 GN=Chil3 PE=1 SV=2 | sp\|O35744\|CHIL3_MOUSE | 5.02E+07 |
| 11 | Endoplasmic reticulum chaperone BiP OS=Mus musculus OX=10090 GN=Hspa5 PE=1 SV=3 | sp\|P20029\|BIP_MOUSE (+5) | 4.95E+07 |
| 12 | High mobility group protein B2 OS=Mus musculus GN=Hmgb2 PE=1 SV=3 | sp\|P30681\|HMGB2_MOUSE (+3) | 4.93E+07 |
| 13 | Peroxiredoxin-2 OS=Mus musculus GN=Prdx2 PE=1 SV=3 | sp\|Q61171\|PRDX2_MOUSE | 4.83E+07 |
| 14 | Superoxide dismutase [Cu-Zn] OS=Mus musculus GN=Sod1 PE=1 SV=2 | sp\|P08228\|SODC_MOUSE | 4.82E+07 |
| 15 | Heat shock protein HSP 90-beta OS=Mus musculus GN=Hsp90ab1 PE=1 SV=3 | sp\|P11499\|HS90B_MOUSE | 4.68E+07 |
| 16 | Heat shock protein HSP 90-alpha OS=Mus musculus GN=Hsp90aa1 PE=1 SV=4 | sp\|P07901\|HS90A_MOUSE (+1) | 4.68E+07 |
| 17 | Histone H2A.J OS=Mus musculus GN=H2afj PE=1 SV=1 | sp\|Q8R1M2\|H2AJ_MOUSE | 4.37E+07 |
| 18 | Histone H2B type 3-B OS=Mus musculus GN=Hist3h2bb PE=1 SV=3 | sp\|Q8CGP0\|H2B3B_MOUSE (+1) | 4.32E+07 |
| 19 | Nucleophosmin OS=Mus musculus GN=Npm1 PE=1 SV=1 | sp\|Q61937\|NPM_MOUSE (+2) | 4.23E+07 |
| 20 | Calreticulin OS=Mus musculus GN=Calr PE=1 SV=1 | sp\|P14211\|CALR_MOUSE (+1) | 4.23E+07 |

| | | | |
|---|---|---|---|
| 21 | Peptidyl-prolyl cis-trans isomerase A OS=Mus musculus GN=Ppia PE=1 SV=2 | sp\|P17742\|PPIA_MOUSE | 4.12E+07 |
| 22 | Histone H4 OS=Mus musculus GN=Hist1h4a PE=1 SV=2 | sp\|P62806\|H4_MOUSE | 4.09E+07 |
| 23 | Profilin-1 OS=Mus musculus GN=Pfn1 PE=1 SV=2 | sp\|P62962\|PROF1_MOUSE | 3.81E+07 |
| 24 | Isoform 2 of Heterogeneous nuclear ribonucleoprotein A3 OS=Mus musculus OX=10090 GN=Hnrnpa3 | sp\|Q8BG05-2\|ROA3_MOUSE (+2) | 3.30E+07 |
| 25 | Transketolase OS=Mus musculus GN=Tkt PE=1 SV=1 | sp\|P40142\|TKT_MOUSE | 3.26E+07 |
| 26 | Histone H1.2 OS=Mus musculus GN=Hist1h1c PE=1 SV=2 | sp\|P15864\|H12_MOUSE | 3.25E+07 |
| 27 | Ferritin heavy chain OS=Mus musculus GN=Fth1 PE=1 SV=2 | sp\|P09528\|FRIH_MOUSE | 3.12E+07 |
| 28 | Alpha-hemoglobin-stabilizing protein OS=Mus musculus OX=10090 GN=Ahsp PE=1 SV=1 | sp\|Q9CY02\|AHSP_MOUSE | 3.05E+07 |
| 29 | High mobility group protein B1 OS=Mus musculus GN=Hmgb1 PE=1 SV=2 | sp\|P63158\|HMGB1_MOUSE (+8) | 3.01E+07 |
| 30 | Elongation factor 2 OS=Mus musculus GN=Eef2 PE=1 SV=2 | sp\|P58252\|EF2_MOUSE | 2.95E+07 |
| 31 | Neutrophilic granule protein OS=Mus musculus OX=10090 GN=Ngp PE=1 SV=1 | sp\|O08692\|NGP_MOUSE | 2.92E+07 |
| 32 | Protein S100-A8 OS=Mus musculus OX=10090 GN=S100a8 PE=1 SV=3 | sp\|P27005\|S10A8_MOUSE | 2.84E+07 |
| 33 | Glyceraldehyde-3-phosphate dehydrogenase OS=Mus musculus GN=Gapdh PE=1 SV=2 | sp\|P16858\|G3P_MOUSE (+2) | 2.82E+07 |
| 34 | Tubulin beta-5 chain OS=Mus musculus GN=Tubb5 PE=1 SV=1 | sp\|P99024\|TBB5_MOUSE | 2.81E+07 |
| 35 | Tubulin beta-4B chain OS=Mus musculus GN=Tubb4b PE=1 SV=1 | sp\|P68372\|TBB4B_MOUSE | 2.81E+07 |
| 36 | Tubulin alpha-1B chain OS=Mus musculus GN=Tuba1b PE=1 SV=2 | sp\|P05213\|TBA1B_MOUSE | 2.79E+07 |
| 37 | Tubulin alpha-4A chain OS=Mus musculus GN=Tuba4a PE=1 SV=1 | sp\|P68368\|TBA4A_MOUSE (+1) | 2.79E+07 |
| 38 | Eukaryotic translation initiation factor 5A-1 OS=Mus musculus GN=Eif5a PE=1 SV=2 | sp\|P63242\|IF5A1_MOUSE (+1) | 2.75E+07 |
| 39 | Isoform 2 of Heterogeneous nuclear ribonucleoproteins A2/B1 OS=Mus musculus OX=10090 GN=Hnrnpa2b1 | sp\|O88569-2\|ROA2_MOUSE (+3) | 2.71E+07 |
| 40 | Lactotransferrin OS=Mus musculus GN=Ltf PE=1 SV=4 | sp\|P08071\|TRFL_MOUSE (+2) | 2.61E+07 |
| 41 | Small ribosomal subunit protein eS12 OS=Mus musculus OX=10090 GN=Rps12 PE=1 SV=3 | sp\|P63323\|RS12_MOUSE | 2.38E+07 |

| | | | |
|---|---|---|---|
| 42 | Histone H3.1 OS=Mus musculus GN=Hist1h3a PE=1 SV=2 | sp\|P68433\|H31_MOUSE (+4) | 2.31E+07 |
| 43 | 60S acidic ribosomal protein P2 OS=Mus musculus GN=Rplp2 PE=1 SV=3 | sp\|P99027\|RLA2_MOUSE | 2.20E+07 |
| 44 | Heterogeneous nuclear ribonucleoprotein A1 OS=Mus musculus GN=Hnrnpa1 PE=1 SV=2 | sp\|P49312\|ROA1_MOUSE (+4) | 2.15E+07 |
| 45 | RNA-binding protein FUS OS=Mus musculus GN=Fus PE=2 SV=1 | sp\|P56959\|FUS_MOUSE (+6) | 2.10E+07 |
| 46 | Malate dehydrogenase, mitochondrial OS=Mus musculus GN=Mdh2 PE=1 SV=3 | sp\|P08249\|MDHM_MOUSE | 2.03E+07 |
| 47 | Transaldolase OS=Mus musculus GN=Taldo1 PE=1 SV=2 | sp\|Q93092\|TALDO_MOUSE (+1) | 2.02E+07 |
| 48 | Uncharacterized protein OS=Mus musculus OX=10090 GN=Psap PE=2 SV=1 | tr\|Q3UE29\|Q3UE29_MOUSE | 1.96E+07 |
| 49 | Isoform 2 of Immunoglobulin heavy constant mu OS=Mus musculus OX=10090 GN=Ighm | sp\|P01872-2\|IGHM_MOUSE (+3) | 1.92E+07 |
| 50 | 40S ribosomal protein S20 OS=Mus musculus GN=Rps20 PE=1 SV=1 | sp\|P60867\|RS20_MOUSE | 1.91E+07 |
| 51 | Cofilin-1 OS=Mus musculus GN=Cfl1 PE=1 SV=3 | sp\|P18760\|COF1_MOUSE | 1.90E+07 |
| 52 | Ezrin OS=Mus musculus GN=Ezr PE=1 SV=3 | sp\|P26040\|EZRI_MOUSE (+3) | 1.87E+07 |
| 53 | Stathmin OS=Mus musculus GN=Stmn1 PE=1 SV=2 | sp\|P54227\|STMN1_MOUSE (+7) | 1.84E+07 |
| 54 | Nucleoside diphosphate kinase OS=Mus musculus OX=10090 GN=Gm20390 PE=3 SV=1 | tr\|E9PZF0\|E9PZF0_MOUSE | 1.80E+07 |
| 55 | Peroxiredoxin-1 OS=Mus musculus GN=Prdx1 PE=1 SV=1 | sp\|P35700\|PRDX1_MOUSE (+1) | 1.77E+07 |
| 56 | GTP-binding nuclear protein Ran OS=Mus musculus GN=Ran PE=1 SV=3 | sp\|P62827\|RAN_MOUSE (+1) | 1.75E+07 |
| 57 | RNA-binding protein 3 OS=Mus musculus GN=Rbm3 PE=1 SV=1 | sp\|O89086\|RBM3_MOUSE | 1.74E+07 |
| 58 | Endoplasmin OS=Mus musculus GN=Hsp90b1 PE=1 SV=2 | sp\|P08113\|ENPL_MOUSE (+2) | 1.74E+07 |
| 59 | 40S ribosomal protein S3a OS=Mus musculus GN=Rps3a PE=1 SV=3 | sp\|P97351\|RS3A_MOUSE (+3) | 1.74E+07 |
| 60 | 60 kDa heat shock protein, mitochondrial OS=Mus musculus GN=Hspd1 PE=1 SV=1 | sp\|P63038\|CH60_MOUSE | 1.73E+07 |
| 61 | 60S ribosomal protein L7 OS=Mus musculus GN=Rpl7 PE=1 SV=2 | sp\|P14148\|RL7_MOUSE (+1) | 1.72E+07 |
| 62 | Isoform 2 of Acidic leucine-rich nuclear phosphoprotein 32 family member B OS=Mus musculus OX=10090 GN=Anp32b | sp\|Q9EST5-2\|AN32B_MOUSE (+2) | 1.65E+07 |
| 63 | Alpha-enolase OS=Mus musculus GN=Eno1 PE=1 SV=3 | sp\|P17182\|ENOA_MOUSE | 1.65E+07 |

| | | | |
|---|---|---|---|
| 64 | Thioredoxin OS=Mus musculus GN=Txn PE=1 SV=3 | sp\|P10639\|THIO_MOUSE | 1.64E+07 |
| 65 | L-lactate dehydrogenase A chain OS=Mus musculus GN=Ldha PE=1 SV=3 | sp\|P06151\|LDHA_MOUSE (+4) | 1.56E+07 |
| 66 | Heterogeneous nuclear ribonucleoprotein A/B OS=Mus musculus GN=Hnrnpab PE=1 SV=1 | sp\|Q99020\|ROAA_MOUSE (+4) | 1.55E+07 |
| 67 | 14-3-3 protein zeta/delta OS=Mus musculus GN=Ywhaz PE=1 SV=1 | sp\|P63101\|1433Z_MOUSE | 1.51E+07 |
| 68 | Hemogen OS=Mus musculus OX=10090 GN=Hemgn PE=1 SV=1 | sp\|Q9ERZ0\|HEMGN_MOUSE | 1.48E+07 |
| 69 | Isoform Short of 14-3-3 protein beta/alpha OS=Mus musculus OX=10090 GN=Ywhab | sp\|Q9CQV8-2\|1433B_MOUSE (+1) | 1.48E+07 |
| 70 | 40S ribosomal protein S21 OS=Mus musculus GN=Rps21 PE=1 SV=1 | sp\|Q9CQR2\|RS21_MOUSE (+1) | 1.46E+07 |
| 71 | Isoform 2 of Porphobilinogen deaminase OS=Mus musculus OX=10090 GN=Hmbs | sp\|P22907-2\|HEM3_MOUSE (+3) | 1.45E+07 |
| 72 | Serine/arginine-rich splicing factor 2 OS=Mus musculus GN=Srsf2 PE=1 SV=4 | sp\|Q62093\|SRSF2_MOUSE | 1.45E+07 |
| 73 | Proliferating cell nuclear antigen OS=Mus musculus GN=Pcna PE=1 SV=2 | sp\|P17918\|PCNA_MOUSE (+1) | 1.43E+07 |
| 74 | Eukaryotic initiation factor 4A-I OS=Mus musculus GN=Eif4a1 PE=1 SV=1 | sp\|P60843\|IF4A1_MOUSE (+5) | 1.43E+07 |
| 75 | Nucleolin OS=Mus musculus GN=Ncl PE=1 SV=2 | sp\|P09405\|NUCL_MOUSE (+3) | 1.39E+07 |
| 76 | 60S ribosomal protein L18 OS=Mus musculus GN=Rpl18 PE=1 SV=3 | sp\|P35980\|RL18_MOUSE (+6) | 1.38E+07 |
| 77 | Calmodulin-2 OS=Mus musculus OX=10090 GN=Calm2 PE=1 SV=1 | sp\|P0DP27\|CALM2_MOUSE | 1.37E+07 |
| 78 | Polyubiquitin-B OS=Mus musculus GN=Ubb PE=2 SV=1 | sp\|P0CG49\|UBB_MOUSE (+14) | 1.34E+07 |
| 79 | Protein disulfide-isomerase A3 OS=Mus musculus GN=Pdia3 PE=1 SV=2 | sp\|P27773\|PDIA3_MOUSE | 1.34E+07 |
| 80 | Proliferation-associated protein 2G4 OS=Mus musculus GN=Pa2g4 PE=1 SV=3 | sp\|P50580\|PA2G4_MOUSE | 1.32E+07 |
| 81 | 60S ribosomal protein L6 OS=Mus musculus GN=Rpl6 PE=1 SV=3 | sp\|P47911\|RL6_MOUSE (+1) | 1.27E+07 |
| 82 | Ran-specific GTPase-activating protein OS=Mus musculus GN=Ranbp1 PE=1 SV=2 | sp\|P34022\|RANG_MOUSE (+1) | 1.27E+07 |
| 83 | Myeloperoxidase OS=Mus musculus GN=Mpo PE=1 SV=2 | sp\|P11247\|PERM_MOUSE (+3) | 1.26E+07 |
| 84 | 60S ribosomal protein L5 OS=Mus musculus GN=Rpl5 PE=1 SV=3 | sp\|P47962\|RL5_MOUSE (+4) | 1.24E+07 |
| 85 | Receptor of activated protein C kinase 1 OS=Mus musculus GN=Rack1 PE=1 SV=3 | sp\|P68040\|RACK1_MOUSE | 1.20E+07 |

| | | | |
|---|---|---|---|
| 86 | Protein RCC2 OS=Mus musculus GN=Rcc2 PE=1 SV=1 | sp\|Q8BK67\|RCC2_MOUSE | 1.19E+07 |
| 87 | Isoform Smooth muscle of Myosin light polypeptide 6 OS=Mus musculus OX=10090 GN=Myl6 | sp\|Q60605-2\|MYL6_MOUSE (+4) | 1.18E+07 |
| 88 | 60S ribosomal protein L10a OS=Mus musculus GN=Rpl10a PE=1 SV=3 | sp\|P53026\|RL10A_MOUSE (+4) | 1.18E+07 |
| 89 | Hepatoma-derived growth factor OS=Mus musculus GN=Hdgf PE=1 SV=2 | sp\|P51859\|HDGF_MOUSE | 1.15E+07 |
| 90 | Isoform 3 of Heterogeneous nuclear ribonucleoprotein D0 OS=Mus musculus OX=10090 GN=Hnrnpd | sp\|Q60668-3\|HNRPD_MOUSE (+1) | 1.14E+07 |
| 91 | ATP synthase subunit beta, mitochondrial OS=Mus musculus GN=Atp5b PE=1 SV=2 | sp\|P56480\|ATPB_MOUSE | 1.14E+07 |
| 92 | 40S ribosomal protein S28 OS=Mus musculus GN=Rps28 PE=1 SV=1 | sp\|P62858\|RS28_MOUSE (+1) | 1.14E+07 |
| 93 | RNA binding motif protein, X-linked-like-1 OS=Mus musculus GN=Rbmxl1 PE=1 SV=1 | sp\|Q91VM5\|RMXL1_MOUSE (+4) | 1.12E+07 |
| 94 | 60S ribosomal protein L15 OS=Mus musculus GN=Rpl15 PE=2 SV=4 | sp\|Q9CZM2\|RL15_MOUSE (+2) | 1.11E+07 |
| 95 | Annexin A1 OS=Mus musculus GN=Anxa1 PE=1 SV=2 | sp\|P10107\|ANXA1_MOUSE | 1.11E+07 |
| 96 | Isoform 2 of Heterogeneous nuclear ribonucleoprotein K OS=Mus musculus OX=10090 GN=Hnrnpk | sp\|P61979-2\|HNRPK_MOUSE (+5) | 1.09E+07 |
| 97 | Coronin-1A OS=Mus musculus GN=Coro1a PE=1 SV=5 | sp\|O89053\|COR1A_MOUSE (+2) | 1.07E+07 |
| 98 | Isoform 2 of Heterogeneous nuclear ribonucleoprotein U OS=Mus musculus OX=10090 GN=Hnrnpu | sp\|Q8VEK3-2\|HNRPU_MOUSE (+7) | 1.07E+07 |
| 99 | Fructose-bisphosphate aldolase A OS=Mus musculus GN=Aldoa PE=1 SV=2 | sp\|P05064\|ALDOA_MOUSE (+2) | 1.06E+07 |
| 100 | Resistin-like gamma OS=Mus musculus OX=10090 GN=Retnlg PE=1 SV=1 | sp\|Q8K426\|RETNG_MOUSE | 1.05E+07 |
| 101 | Neutrophil gelatinase-associated lipocalin OS=Mus musculus GN=Lcn2 PE=1 SV=1 | sp\|P11672\|NGAL_MOUSE (+2) | 1.04E+07 |
| 102 | Isoform 2 of Protein SET OS=Mus musculus OX=10090 GN=Set | sp\|Q9EQU5-2\|SET_MOUSE (+3) | 1.04E+07 |
| 103 | 60S ribosomal protein L7a OS=Mus musculus GN=Rpl7a PE=1 SV=2 | sp\|P12970\|RL7A_MOUSE (+3) | 1.02E+07 |
| 104 | 14-3-3 protein epsilon OS=Mus musculus GN=Ywhae PE=1 SV=1 | sp\|P62259\|1433E_MOUSE | 1.01E+07 |

| | | | |
|---|---|---|---|
| 105 | Isoform 2 of Elongation factor 1-delta OS=Mus musculus OX=10090 GN=Eef1d | sp\|P57776-2\|EF1D_MOUSE (+2) | 1.00E+07 |
| 106 | Protein disulfide-isomerase OS=Mus musculus GN=P4hb PE=1 SV=2 | sp\|P09103\|PDIA1_MOUSE (+7) | 9834600 |
| 107 | Adenosylhomocysteinase OS=Mus musculus GN=Ahcy PE=1 SV=3 | sp\|P50247\|SAHH_MOUSE (+4) | 9448000 |
| 108 | Oxygen-dependent coproporphyrinogen-III oxidase, mitochondrial OS=Mus musculus GN=Cpox PE=1 SV=2 | sp\|P36552\|HEM6_MOUSE | 9391800 |
| 109 | Isoform Short of Serine/arginine-rich splicing factor 3 OS=Mus musculus OX=10090 GN=Srsf3 | sp\|P84104-2\|SRSF3_MOUSE (+2) | 9115900 |
| 110 | ATP synthase subunit alpha, mitochondrial OS=Mus musculus GN=Atp5a1 PE=1 SV=1 | sp\|Q03265\|ATPA_MOUSE (+1) | 8919000 |
| 111 | 40S ribosomal protein SA OS=Mus musculus GN=Rpsa PE=1 SV=4 | sp\|P14206\|RSSA_MOUSE | 8720700 |
| 112 | 14-3-3 protein eta OS=Mus musculus GN=Ywhah PE=1 SV=2 | sp\|P68510\|1433F_MOUSE | 8674100 |
| 113 | Purine nucleoside phosphorylase OS=Mus musculus GN=Pnp PE=1 SV=2 | sp\|P23492\|PNPH_MOUSE (+1) | 8526300 |
| 114 | Isoform C1 of Heterogeneous nuclear ribonucleoproteins C1/C2 OS=Mus musculus OX=10090 GN=Hnrnpc | sp\|Q9Z204-2\|HNRPC_MOUSE (+5) | 8489400 |
| 115 | Histone-binding protein RBBP4 OS=Mus musculus GN=Rbbp4 PE=1 SV=5 | sp\|Q60972\|RBBP4_MOUSE | 8207800 |
| 116 | 10 kDa heat shock protein, mitochondrial OS=Mus musculus GN=Hspe1 PE=1 SV=2 | sp\|Q64433\|CH10_MOUSE | 7971800 |
| 117 | 40S ribosomal protein S6 OS=Mus musculus GN=Rps6 PE=1 SV=1 | sp\|P62754\|RS6_MOUSE (+2) | 7917000 |
| 118 | 60S ribosomal protein L23a OS=Mus musculus GN=Rpl23a PE=1 SV=1 | sp\|P62751\|RL23A_MOUSE (+2) | 7805100 |
| 119 | Splicing factor, proline- and glutamine-rich OS=Mus musculus GN=Sfpq PE=1 SV=1 | sp\|Q8VIJ6\|SFPQ_MOUSE | 7708900 |
| 120 | 60S ribosomal protein L12 OS=Mus musculus GN=Rpl12 PE=1 SV=2 | sp\|P35979\|RL12_MOUSE (+2) | 7676800 |
| 121 | Peroxiredoxin-6 OS=Mus musculus GN=Prdx6 PE=1 SV=3 | sp\|O08709\|PRDX6_MOUSE | 7591700 |
| 122 | Transitional endoplasmic reticulum ATPase OS=Mus musculus GN=Vcp PE=1 SV=4 | sp\|Q01853\|TERA_MOUSE (+1) | 7080400 |
| 123 | Far upstream element-binding protein 2 OS=Mus musculus GN=Khsrp PE=1 SV=2 | sp\|Q3U0V1\|FUBP2_MOUSE (+1) | 7002700 |
| 124 | 40S ribosomal protein S19 OS=Mus musculus GN=Rps19 PE=1 SV=3 | sp\|Q9CZX8\|RS19_MOUSE (+5) | 7000900 |

| | | | |
|---|---|---|---|
| 125 | 40S ribosomal protein S8 OS=Mus musculus GN=Rps8 PE=1 SV=2 | sp\|P62242\|RS8_MOUSE (+1) | 6965000 |
| 126 | Aconitate hydratase, mitochondrial OS=Mus musculus GN=Aco2 PE=1 SV=1 | sp\|Q99KI0\|ACON_MOUSE | 6926000 |
| 127 | Elongation factor 1-beta OS=Mus musculus GN=Eef1b PE=1 SV=5 | sp\|O70251\|EF1B_MOUSE (+1) | 6689300 |
| 128 | Glucose-6-phosphate isomerase OS=Mus musculus GN=Gpi PE=1 SV=4 | sp\|P06745\|G6PI_MOUSE | 6543100 |
| 129 | Stress-induced-phosphoprotein 1 OS=Mus musculus GN=Stip1 PE=1 SV=1 | sp\|Q60864\|STIP1_MOUSE (+1) | 6472800 |
| 130 | Isoform 2 of Transcription factor BTF3 OS=Mus musculus OX=10090 GN=Btf3 | sp\|Q64152-2\|BTF3_MOUSE | 6461100 |
| 131 | C-1-tetrahydrofolate synthase, cytoplasmic OS=Mus musculus GN=Mthfd1 PE=1 SV=4 | sp\|Q922D8\|C1TC_MOUSE (+3) | 6165000 |
| 132 | 40S ribosomal protein S3 OS=Mus musculus GN=Rps3 PE=1 SV=1 | sp\|P62908\|RS3_MOUSE (+1) | 6140700 |
| 133 | Alpha-actinin-4 OS=Mus musculus GN=Actn4 PE=1 SV=1 | sp\|P57780\|ACTN4_MOUSE (+3) | 6083100 |
| 134 | Serine/arginine-rich splicing factor 1 OS=Mus musculus GN=Srsf1 PE=1 SV=3 | sp\|Q6PDM2\|SRSF1_MOUSE (+1) | 6076400 |
| 135 | Heterogeneous nuclear ribonucleoprotein L OS=Mus musculus GN=Hnrnpl PE=1 SV=2 | sp\|Q8R081\|HNRPL_MOUSE (+2) | 6064200 |
| 136 | Citrate synthase, mitochondrial OS=Mus musculus GN=Cs PE=1 SV=1 | sp\|Q9CZU6\|CISY_MOUSE | 6060000 |
| 137 | Rho GDP-dissociation inhibitor 1 OS=Mus musculus GN=Arhgdia PE=1 SV=3 | sp\|Q99PT1\|GDIR1_MOUSE | 6034100 |
| 138 | Isoform 2 of Tropomyosin alpha-3 chain OS=Mus musculus OX=10090 GN=Tpm3 | sp\|P21107-2\|TPM3_MOUSE (+5) | 6008900 |
| 139 | Cathelicidin antimicrobial peptide OS=Mus musculus GN=Camp PE=1 SV=2 | sp\|P51437\|CAMP_MOUSE (+3) | 5886700 |
| 140 | 60S acidic ribosomal protein P0 OS=Mus musculus GN=Rplp0 PE=1 SV=3 | sp\|P14869\|RLA0_MOUSE (+2) | 5846200 |
| 141 | MYB-1b OS=Mus musculus OX=10090 GN=Ybx1 PE=2 SV=1 | tr\|Q60951\|Q60951_MOUSE | 5689200 |
| 142 | Triosephosphate isomerase OS=Mus musculus GN=Tpi1 PE=1 SV=4 | sp\|P17751\|TPIS_MOUSE (+1) | 5617000 |
| 143 | Ubiquitin-conjugating enzyme E2 L3 OS=Mus musculus GN=Ube2l3 PE=1 SV=1 | sp\|P68037\|UB2L3_MOUSE (+2) | 5600500 |
| 144 | 40S ribosomal protein S7 OS=Mus musculus GN=Rps7 PE=2 SV=1 | sp\|P62082\|RS7_MOUSE | 5299700 |
| 145 | Filamin-A OS=Mus musculus GN=Flna PE=1 SV=5 | sp\|Q8BTM8\|FLNA_MOUSE (+3) | 5257600 |
| 146 | Myosin-9 OS=Mus musculus GN=Myh9 PE=1 SV=4 | sp\|Q8VDD5\|MYH9_MOUSE | 5237400 |

| | | | |
|---|---|---|---|
| 147 | Lupus La protein homolog OS=Mus musculus GN=Ssb PE=1 SV=1 | sp\|P32067\|LA_MOUSE (+3) | 5174600 |
| 148 | 40S ribosomal protein S2 OS=Mus musculus GN=Rps2 PE=1 SV=3 | sp\|P25444\|RS2_MOUSE (+6) | 5047400 |
| 149 | Isoform 2 of Heterogeneous nuclear ribonucleoprotein F OS=Mus musculus OX=10090 GN=Hnrnpf | sp\|Q9Z2X1-2\|HNRPF_MOUSE (+1) | 4777200 |
| 150 | Non-POU domain-containing octamer-binding protein OS=Mus musculus GN=Nono PE=1 SV=3 | sp\|Q99K48\|NONO_MOUSE (+5) | 4527600 |
| 151 | Peptidyl-prolyl cis-trans isomerase FKBP4 OS=Mus musculus GN=Fkbp4 PE=1 SV=5 | sp\|P30416\|FKBP4_MOUSE (+1) | 4473100 |
| 152 | 60S ribosomal protein L11 OS=Mus musculus GN=Rpl11 PE=1 SV=4 | sp\|Q9CXW4\|RL11_MOUSE | 4456400 |
| 153 | Peptidyl-prolyl cis-trans isomerase B OS=Mus musculus GN=Ppib PE=1 SV=2 | sp\|P24369\|PPIB_MOUSE | 4450700 |
| 154 | Nascent polypeptide-associated complex subunit alpha, muscle-specific form OS=Mus musculus GN=Naca PE=1 SV=2 | sp\|P70670\|NACAM_MOUSE | 4409900 |
| 155 | T-complex protein 1 subunit theta OS=Mus musculus GN=Cct8 PE=1 SV=3 | sp\|P42932\|TCPQ_MOUSE (+6) | 4347200 |
| 156 | Nuclear autoantigenic sperm protein OS=Mus musculus GN=Nasp PE=1 SV=2 | sp\|Q99MD9\|NASP_MOUSE (+2) | 4205600 |
| 157 | Uroporphyrinogen decarboxylase OS=Mus musculus GN=Urod PE=1 SV=2 | sp\|P70697\|DCUP_MOUSE | 4084300 |
| 158 | Clathrin heavy chain 1 OS=Mus musculus GN=Cltc PE=1 SV=3 | sp\|Q68FD5\|CLH1_MOUSE (+3) | 4015200 |
| 159 | Isoform Cytoplasmic+peroxisomal of Peroxiredoxin-5, mitochondrial OS=Mus musculus OX=10090 GN=Prdx5 | sp\|P99029-2\|PRDX5_MOUSE (+6) | 4012100 |
| 160 | Vimentin OS=Mus musculus GN=Vim PE=1 SV=3 | sp\|P20152\|VIME_MOUSE (+6) | 3798600 |
| 161 | Isoform 2 of Calpastatin OS=Mus musculus OX=10090 GN=Cast | sp\|P51125-2\|ICAL_MOUSE (+10) | 3662200 |
| 162 | Cysteine and histidine-rich domain-containing protein 1 OS=Mus musculus GN=Chordc1 PE=1 SV=1 | sp\|Q9D1P4\|CHRD1_MOUSE (+1) | 3655200 |
| 163 | DNA replication licensing factor MCM7 OS=Mus musculus GN=Mcm7 PE=1 SV=1 | sp\|Q61881\|MCM7_MOUSE (+2) | 3441800 |
| 164 | Igk protein OS=Mus musculus OX=10090 GN=Igk PE=1 SV=1 | tr\|I6L978\|I6L978_MOUSE | 3230800 |

| | | | |
|---|---|---|---|
| 165 | Talin-1 OS=Mus musculus GN=Tln1 PE=1 SV=2 | sp\|P26039\|TLN1_MOUSE (+2) | 3141400 |
| 166 | Isoform 2 of Acidic leucine-rich nuclear phosphoprotein 32 family member E OS=Mus musculus OX=10090 GN=Anp32e | sp\|P97822-2\|AN32E_MOUSE (+3) | 3135700 |
| 167 | Plasminogen activator inhibitor 1 RNA-binding protein OS=Mus musculus GN=Serbp1 PE=1 SV=2 | sp\|Q9CY58\|PAIRB_MOUSE | 3064700 |
| 168 | Delta-aminolevulinic acid dehydratase OS=Mus musculus GN=Alad PE=1 SV=1 | sp\|P10518\|HEM2_MOUSE (+1) | 3016900 |
| 169 | Parkinson disease protein 7 homolog OS=Mus musculus OX=10090 GN=Park7 PE=1 SV=1 | sp\|Q99LX0\|PARK7_MOUSE (+5) | 2944700 |
| 170 | Malate dehydrogenase, cytoplasmic OS=Mus musculus GN=Mdh1 PE=1 SV=3 | sp\|P14152\|MDHC_MOUSE (+1) | 2924700 |
| 171 | Phosphatidylethanolamine-binding protein 1 OS=Mus musculus GN=Pebp1 PE=1 SV=3 | sp\|P70296\|PEBP1_MOUSE (+1) | 2922100 |
| 172 | Protein TANC2 OS=Mus musculus GN=Tanc2 PE=1 SV=1 | sp\|A2A690\|TANC2_MOUSE | 2770400 |
| 173 | DNA replication licensing factor MCM4 OS=Mus musculus GN=Mcm4 PE=1 SV=1 | sp\|P49717\|MCM4_MOUSE (+5) | 2710500 |
| 174 | Protein S100-A11 OS=Mus musculus GN=S100a11 PE=1 SV=1 | sp\|P50543\|S10AB_MOUSE | 2309000 |
| 175 | Isoform 2 of Rab GDP dissociation inhibitor beta OS=Mus musculus OX=10090 GN=Gdi2 | sp\|Q61598-2\|GDIB_MOUSE (+5) | 2244800 |
| 176 | DNA replication licensing factor MCM3 OS=Mus musculus GN=Mcm3 PE=1 SV=2 | sp\|P25206\|MCM3_MOUSE | 2126800 |
| 177 | 28 kDa heat- and acid-stable phosphoprotein OS=Mus musculus GN=Pdap1 PE=1 SV=1 | sp\|Q3UHX2\|HAP28_MOUSE | 2074500 |
| 178 | T-complex protein 1 subunit zeta OS=Mus musculus GN=Cct6a PE=1 SV=3 | sp\|P80317\|TCPZ_MOUSE (+5) | 1897800 |
| 179 | Isoform CW17E of Splicing factor 1 OS=Mus musculus OX=10090 GN=Sf1 | sp\|Q64213-2\|SF01_MOUSE (+6) | 1830700 |
| 180 | RIKEN cDNA 2210010C04 gene OS=Mus musculus OX=10090 GN=2210010C04Rik PE=1 SV=1 | tr\|Q9CPN9\|Q9CPN9_MOUSE (+1) | 1600400 |
| 181 | Plastin-2 OS=Mus musculus GN=Lcp1 PE=1 SV=4 | sp\|Q61233\|PLSL_MOUSE (+2) | 1257400 |
| 182 | 6-phosphogluconate dehydrogenase, decarboxylating OS=Mus musculus GN=Pgd PE=1 SV=3 | sp\|Q9DCD0\|6PGD_MOUSE | 1233000 |
| 183 | 60S ribosomal protein L4 OS=Mus musculus GN=Rpl4 PE=1 SV=3 | sp\|Q9D8E6\|RL4_MOUSE (+1) | 1207500 |
| 184 | Ubiquitin-like modifier-activating enzyme 1 OS=Mus musculus GN=Uba1 PE=1 SV=1 | sp\|Q02053\|UBA1_MOUSE (+1) | 1173400 |

| | | | |
|---|---|---|---|
| 185 | Splicing factor U2AF 35 kDa subunit OS=Mus musculus GN=U2af1 PE=1 SV=4 | sp\|Q9D883\|U2AF1_MOUSE | 1032400 |
| 186 | T-complex protein 1 subunit beta OS=Mus musculus GN=Cct2 PE=1 SV=4 | sp\|P80314\|TCPB_MOUSE | 749550 |
| 187 | PC4 and SFRS1-interacting protein OS=Mus musculus GN=Psip1 PE=1 SV=1 | sp\|Q99JF8\|PSIP1_MOUSE | 206570 |
| 188 | Isoform 1 of RNA-binding protein Raly OS=Mus musculus OX=10090 GN=Raly | sp\|Q64012-2\|RALY_MOUSE (+3) | 112230 |
| 189 | 40S ribosomal protein S15 OS=Mus musculus GN=Rps15 PE=2 SV=2 | sp\|P62843\|RS15_MOUSE (+3) | 41277 |
| 190 | Isoform 2 of Calmodulin-regulated spectrin-associated protein 2 OS=Mus musculus OX=10090 GN=Camsap2 | sp\|Q8C1B1-2\|CAMP2_MOUSE (+5) | 4,966.30 |
